# The multivalency of the glucocorticoid receptor ligand-binding domain explains its manifold physiological activities

**DOI:** 10.1101/2021.10.01.462734

**Authors:** Alba Jiménez-Panizo, Andrea Alegre-Martí, Gregory Fettweis, Montserrat Abella, Rosa Antón, Theophilus Tettey, Louis R. Schiltz, Thomas A Johnson, Israel Nuñez-Barrios, Joan Font-Díaz, Carme Caelles, Annabel F. Valledor, Paloma Pérez, Ana M. Rojas, Juan Fernández-Recio, Diego M. Presman, Gordon L. Hager, Pablo Fuentes-Prior, Eva Estébanez-Perpiñá

**Affiliations:** Structural Biology of Nuclear Receptors, Department of Biochemistry and Molecular Biomedicine, Faculty of Biology, University of Barcelona (UB), 08028 Barcelona, Spain; Institute of Biomedicine of the University of Barcelona (IBUB), University of Barcelona (UB), 08028 Barcelona, Spain; Receptor Biology and Gene Expression, National Cancer Institute, National Institutes of Health, Bethesda, Maryland 20892-5055, USA; Molecular Bases of Disease, Biomedical Research Institute Sant Pau (IIB Sant Pau), 08041 Barcelona, Spain; Computational Biology and Bioinformatics. Andalusian Center for Developmental Biology (CABD-CSIC). Campus Universitario Pablo de Olavide, 41013 Sevilla, Spain; Department of Cell Biology, Physiology and Immunology, Faculty of Biology, University of Barcelona, 08028 Barcelona, Spain; Department of Biochemistry and Physiology, Faculty of Pharmacy and Food Sciences, University of Barcelona, Barcelona 08028, Spain; Instituto de Biomedicina de Valencia (IBV)-CSIC, 46010, Valencia, Spain; Instituto de Ciencias de la Vid y del Vino (ICVV), CSIC - Universidad de La Rioja - Gobierno de La Rioja, 26007 Logroño, Spain; IFIBYNE, UBA-CONICET, Universidad de Buenos Aires, Facultad de Ciencias Exactas y Naturales, Buenos Aires, C1428EGA, Argentina

**Keywords:** glucocorticoid receptor / ligand-binding domain / homodimerization / quaternary structure / X-ray crystallography / fluorescence microscopy

## Abstract

The glucocorticoid receptor (GR) is a ubiquitously expressed transcription factor that controls metabolic and homeostatic processes essential for life. Although numerous crystal structures of the GR ligand-binding domain (GR-LBD) have been reported, the functional oligomeric state of the full-length receptor, which is essential for its transcriptional activity, remains disputed. Here we present five new crystal structures of agonist-bound GR-LBD, along with a thorough analysis of previous structural work. Biologically relevant homodimers were identified by studying a battery of GR point mutants including crosslinking assays in solution and quantitative fluorescence microscopy in living cells. Our results highlight the relevance of non-canonical dimerization modes for GR, especially of contacts made by loop L1-3 residues such as Tyr545. Our work unveils likely pathophysiologically relevant quaternary assemblies of the nuclear receptor with important implications for glucocorticoid action and drug design.

## Introduction

Nuclear receptors (NRs) are a superfamily of transcription factors that control central physiological processes ranging from reproduction and development to metabolism, homeostasis, and ultradian rhythms (Conway-Campbell *et al*, 2012; Busada & Cidlowski, 2017). Steroid receptors form an important subclass of ligand-activated NRs comprising the glucocorticoid receptor (GR/NR3C1) (Fig. 1A), the androgen receptor (AR/NR3C4), the progesterone receptor (PR/NR3C3), the mineralocorticoid receptor (MR/NR3C2), as well as estrogen receptors α and β (ERα/NR3A1 and ERβ/NR3A2, respectively) (Bledsoe et al, 2002; Evans & Mangelsdorf, 2014; Jiménez-Panizo et al, 2019). These proteins share a common modular architecture of a long and unstructured N-terminal domain (NTD) followed by a ‘core’ comprised of a highly conserved DNA-binding domain (DBD), a poorly conserved interdomain linker or hinge, and a moderately conserved C-terminal ligand-binding domain (LBD) (Fig. 1A; see Supplementary Fig. 1 for an alignment of LBD sequences from different species) (Housley *et al*, 1990; Ortlund *et al*, 2007; Meijsing *et al*, 2009). GR binds cholesterol-derived compounds termed glucocorticoids (GCs; either natural compounds such as the main stress hormone, cortisol, or synthetic, e.g., dexamethasone (DEX)) in an internal cavity of the LBD. This ligand-binding pocket (LBP) is allosterically coupled to a solvent-exposed surface responsible for the interaction with coregulators, activation function 2 (AF-2) (Pfaff & Fletterick, 2010; Rogatsky *et al*, 2003). A nearby surface area, topologically equivalent to AR binding function-3 (BF-3) interacts with cochaperones (Estebanez-Perpina *et al*, 2007; Jehle *et al*, 2014). Finally, the LBDs of GR and other related receptors (AR, PR, and MR; referred to as the oxosteroid subfamily) feature a unique C-terminal extension after the last LBD helix (H12), termed F-domain (Jiménez-Panizo *et al*, 2019; Fuentes-Prior *et al*, 2019) (Fig.1B).

**Fig. 1.**
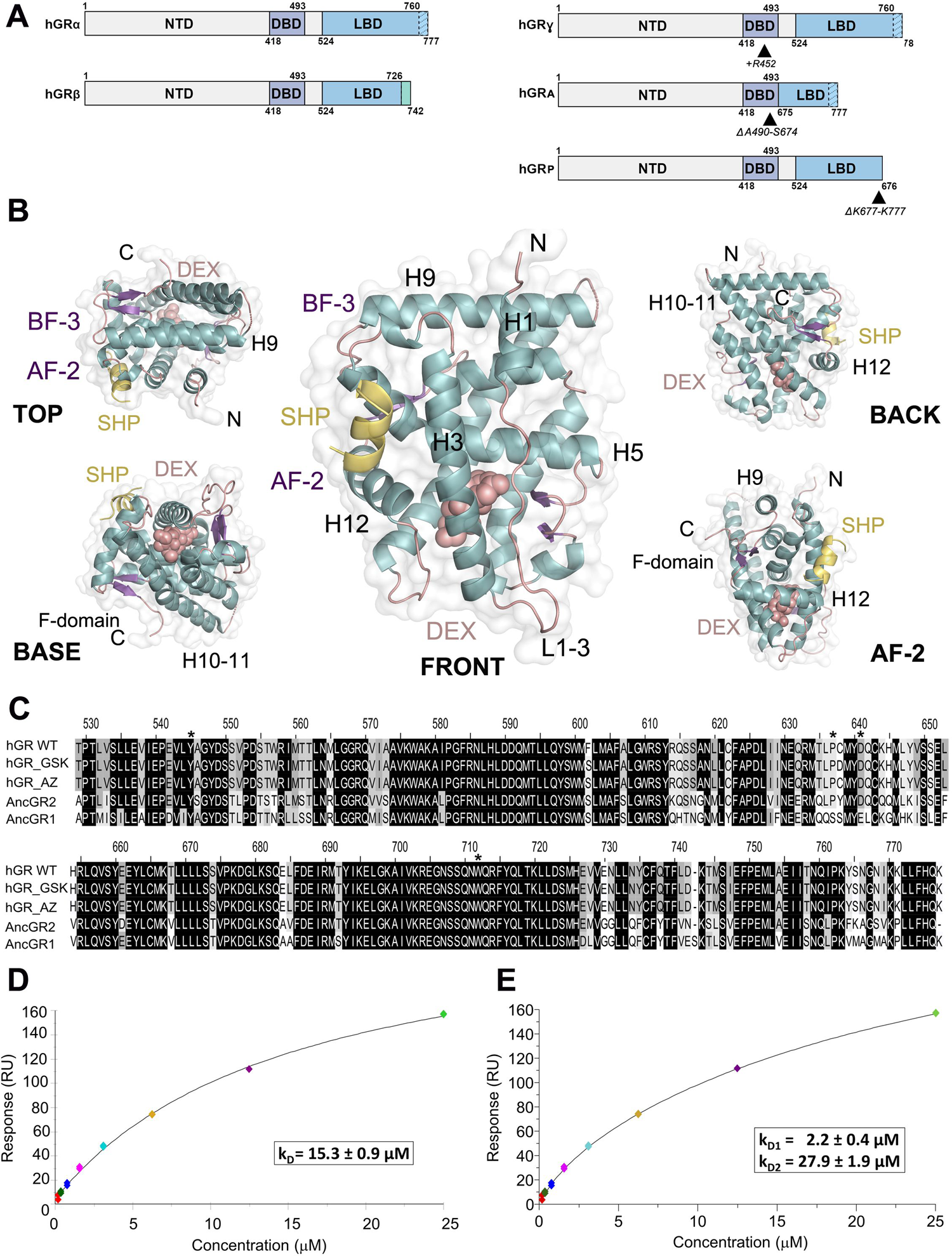
legend: The ligand-binding domain of GR self-associates in solution. **A**, Schematic representation of domain organization in major GR isoforms. GRα and GRβ are identical up to residue 727 (H10/11), but the last 50 (in GRα) and 15 residues (in GRβ, green box) are fully unrelated. Other common isoforms are shown to the right. **B**, Overall structure of the GR-LBD monomer. The domain is shown in standard orientation in the middle of the panel (i.e., with H1 and H3 displayed in the forefront facing the viewer and the AF-2 pocket on the left hand-side of the domain). Four additional orientations are shown to highlight other domain areas. Models are depicted as cartoons with helices (blue), loops (pink) and beta-sheets (purple). The ligand DEX (salmon spheres) and the SHP peptide (yellow cartoon) are also shown. The BF-3 pocket is also labeled. **c**, Sequence alignment of LBDs between wild-type GR, two engineered variants of human GR used in several structure-function investigations (PDB codes 3CLD and 4CSJ), and the resurrected forms, ancGR1 and 2. Strictly conserved residues are white with black shading; other conservatively replaced residues are shaded gray. Residues mutated in the current study are marked with asterisks. **D, E**, SPR analysis of GR-LBD self-association according to (**D**) 1:1 or (**E**) multisite models. The results of experiments conducted in duplicate are shown along with the calculated affinity constants (k_D_). The 1:1 fitting had a Chi^2^ value of 4.52, whereas the multisite model had a Chi^2^ value of 1.25.

*NR3C1* is constitutively expressed in nearly all vertebrate cells. Upon ligand binding, the receptor is trafficked to the nucleus (Vandevyver *et al*, 2012) where a complex DNA-protein interplay modulates its quaternary structure and determines highly dynamic binding to specific chromatin sites (Weikum *et al*, 2017a). Thus, GR integrates signals ranging from available ligands to chromatin remodeling complexes (Clark & Belvisi, 2012) to control a unique set of target genes (up to 17% of the human transcriptome (Franco *et al*, 2019)) to regulate inflammatory responses, cellular proliferation, and differentiation in a highly tissue-specific manner (Oh *et al*, 2017; Cain & Cidlowski, 2017; Sevilla *et al*, 2010). GR also antagonizes the activity of other transcription factors such as activator protein 1 (AP-1) and nuclear factor kB (NF-kB) (De Bosscher *et al*, 2020). Even though monomeric GR is believed to play an important DNA-independent role in the modulation of these major players of the inflammatory response (Louw, 2019), more recent work suggests that both direct DNA binding and GR dimers / tetramers are important for this activity (Presman *et al*, 2014, 2016; Paakinaho *et al*, 2019; Garcia *et al*, 2021; Escoter-Torres *et al*, 2020; Weikum *et al*, 2017c). In line with these manifold functions, alterations in the complex GR signaling pathways due to polymorphisms or mutations in *NR3C1* lead to impaired tissue-specific sensitivity to GCs, which may manifest as either GC resistance (Chrousos syndrome (Chrousos *et al*, 1986)) or hypersensitivity (Nicolaides & Charmandari, 2019). GR is therefore an important pharmacological target to treat several inflammatory pathologies. However, prolonged use or high doses of GCs in patients results in drug resistance and adverse effects (Clark & Belvisi, 2012). Knowledge of GR tertiary and quaternary structures is critical to understand its pivotal functions. Structures of the DBD dimer, both free and DNA-bound have been presented (Frank *et al*, 2018; Hudson *et al*, 2013; Luisi *et al*, 1991; Härd *et al*, 1990), and the LBD has been extensively studied in complex with either agonists or antagonists (Hurt *et al,* 2016; Weikum *et al,* 2017a; Liu *et al*, 2019; Schäcke *et al,* 2007; Biggadike *et al*, 2009; Carson *et al,* 2014; Bledsoe *et al,* 2002).

To date, however, neither full-length (FL) GR nor its core has been structurally characterized. Thus, several important issues regarding the structure-function of GR and related oxosteroid receptors remain unresolved: what is the conformation adopted by dimeric receptors and DNA-bound tetramers *in vivo*, and how do LBD moieties associate in these multimers? Are topologically distinct receptor conformations possible, and are they associated with specific biological functions (e.g., activation vs. repression of transcription)? The answers to these questions have not only an obvious basic science interest, but knowledge of the dimeric/tetrameric conformations of oxosteroid receptors and the detailed mechanism(s) of multimerization would contribute to the design of selective, potent GR modulators that minimize the serious side effects of current drugs.

Here we present a comprehensive structure-and-function investigation of GR multimerization using X-ray crystallography, state-of-the-art bioinformatics tools, surface plasmon resonance (SPR) and crosslinking experiments in solution, and quantitative fluorescence microscopy in live cells. We report five new crystal structures of DEX-bound GR-LBD and integrate this information into the wealth of previous structural data to generate a complete catalog of possible homodimeric arrangements. Four distinct interfaces have been observed to participate in 20 topologically different GR-LBD homodimers. We have identified most favored homodimeric arrangements and suggest how they can combine into pathophysiologically relevant oligomeric assemblies in cells.

## Results

### GR-LBD self-associates in solution

To characterize the ability of GR-LBD to oligomerize in solution, we performed SPR experiments with the ancient variant of the human GR (ancGR2; Fig. 1C). This construct recapitulates the characteristics of wild-type (WT) human GR-LBD and has been repeatedly employed in recent GR structure-function studies because of its higher solubility and stability *in vitro* (Ortlund *et al*, 2007; Weikum *et al*, 2017b). For simplicity, we will refer to all variants of the LBD used for different studies as GR-LBD, unless specific differences are discussed. Briefly, GR-LBD was expressed and purified in the presence of DEX. Agonist-bound GR-LBD was immobilized on CM5 chips using standard amine coupling and increasing concentrations of the same agonist-bound protein (between 0.2 and 25 µM) were run over as analyte.

Although GR-LBD immobilization to the CM5 chip might occlude some protein-protein interaction surfaces, the results of these SPR experiments clearly demonstrate interactions between soluble and immobilized molecules. Several kinetics models were used to interpret the obtained SPR data (Figs. 1D and 1E; representative sensorgrams are shown in Supplementary Fig. 2). GR-LBD self-association behavior could be fitted to a non-covalent, 1:1 Langmuir model with an affinity constant (kD) of 15.3 ± 0.9 µM (Fig. 1D). Interestingly, a significant better fit of the data was achieved using a model of non-covalent multisite interaction, with two independent binding sites (kD1 = 2.2 ± 0.4 µM and kD2 = 27.9 ± 1.9 µM, respectively; Fig. 1E). These results are consistent with GR-LBD tetramer formation, as previously reported with FL-GR in live cells (Presman *et al*, 2016).

### Novel crystal structures of agonist-bound GR-LBD highlight its versatility for self-association

Next, we performed crystallization trials with DEX-bound GR-LBD in the presence of the AF-2 targeting peptide Gln12-Lys30 from the small heterodimer partner (SHP/NR0B2), which contains the canonical LXXLL motif. We conducted solubility screens using all commercially available kits (over 4,800 conditions), which allowed us to identify several new crystallization conditions. Diffraction data from flash-frozen crystals that belong to five different space groups (C2, P3_1_, P6_1_, I4_1_22 and I4_1_32, from lower to higher symmetry) were collected using synchrotron radiation. Major features of the inter-monomer contacts observed in these new GR-LBD structures are briefly summarized below (Fig. 2; see Supplementary Table 1 for a summary of diffraction data, refinement statistics, and model quality).

**Fig. 2.**
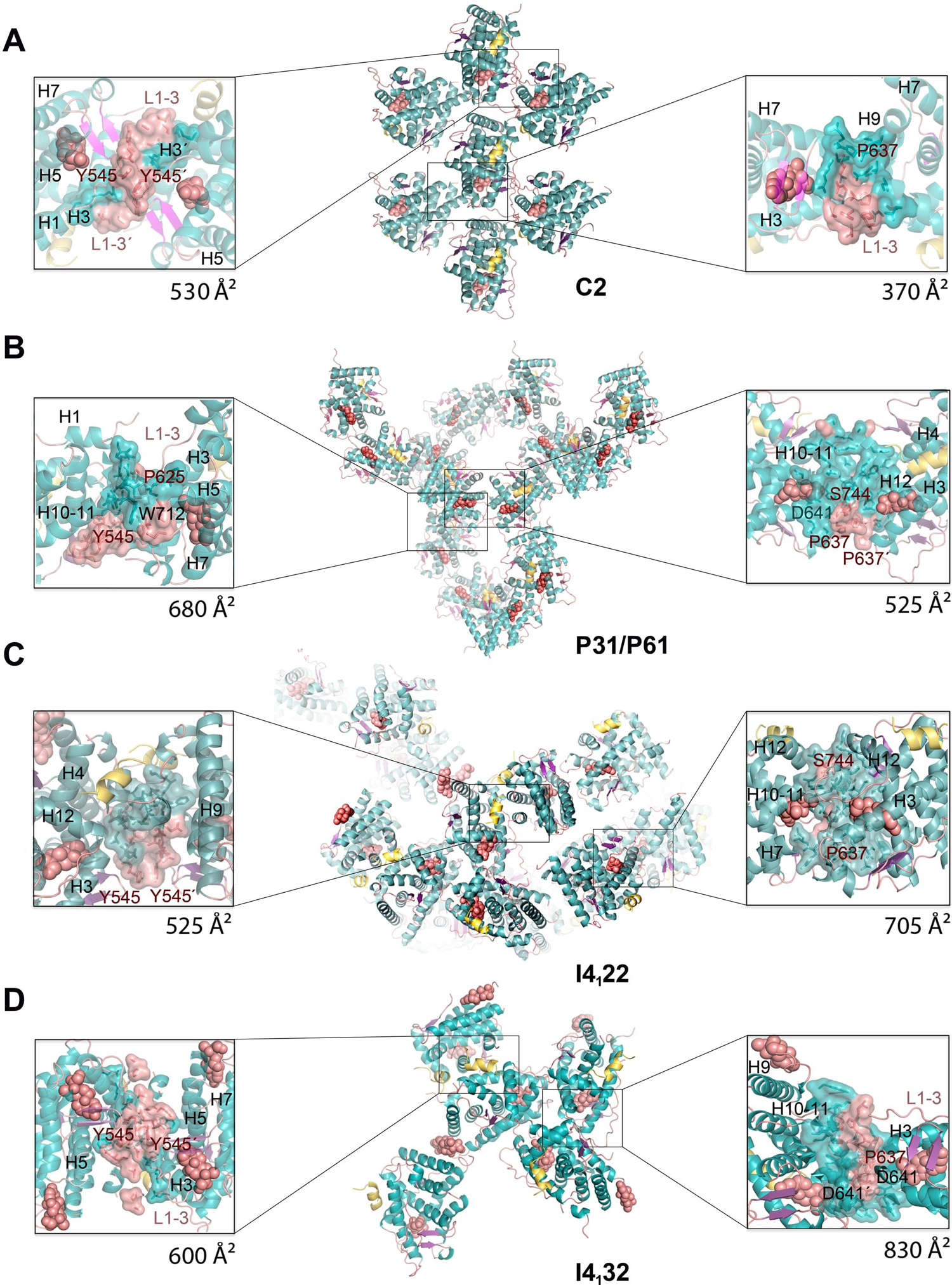
legend: New crystal structures of DEX-bound GR-LBD reveal a variety of quaternary assemblies. For all structures, the overall crystal packing is shown in the central panels. Monomers are depicted as cartoons, DEX molecules are represented as salmon spheres, and SHP peptides as yellow ribbons. Details of intermonomer interfaces are given in the lateral panels, in which the side chains of interacting residues are shown as sticks with all their non-hydrogen atoms. **A**, Monoclinic (C2) crystals. Note that major contacts are centered on L1-3 with stacked Tyr545 phenol rings from two neighboring molecules. **B**, Trigonal (P3_1_) and hexagonal (P6_1_) crystals. The P6_1_ structure generates from the lower symmetry group by conversion of a local (approximate) into a crystallographic (exact) two-fold axis. Residue Tyr545 engages in heterologous contacts with a neighboring molecule in these crystal forms (see the position of the Trp712’ side chain). **C, D**, Common packing of tetragonal (I4_1_22) and cubic (I4_1_32) crystals. Note that the phenol rings of two Tyr545 residues stack as in the C2 crystals, although the two interacting monomers are fully differently oriented relative to each other. Note also that the largest interface in these crystal forms features abutting Asp641 side chains from three monomers, which are organized around a local (I4_1_22) or exact 3-fold axis (I4_1_32; right side panel in **D**).

Crystals of the C2 space group contain a single molecule of GR-LBD·DEX complexed with the SHP peptide, which is well defined by electron density occupying the AF-2 cleft. Two different inter-monomer contacts were identified: the larger, symmetric interaction surface is centered on the L1-3 loops of both monomers and is stabilized by aromatic π-stacking interactions between opposite Tyr545/Tyr545’ residues (Fig. 2A; residues from the second monomer are primed). Further stability is provided by a network of hydrogen bonds (H-bonds) involving several charged residues from both moieties, most notably Asp549 (L1-3), Arg569 (H3), and Asp626 (β-strand S1). A significantly smaller, asymmetric interface features Glu688 (H9), whose carboxylate engages in strong H-bonds with the main chain N atom and the hydroxyl of Ser556’ (L1-3). Additional interactions involve H1 (Leu532) and H9 residues (Lys695, Lys699) from one LBD molecule facing H6 (Glu632’) from the neighbor. Residue Pro637’ (L5-6) is part of this interface, making strong Van der Waals (VdW) interactions with the aliphatic part of Glu688.

Two additional, related crystal structures were solved in the enantiomorphic trigonal and hexagonal space groups, P3_1_ and P6_1_ (Fig. 2B; Supplementary Table 1). In the GR-LBD·DEX homodimer with the larger interface, the cleft between H9 and H10-11 is filled with side chains from neighboring L1-3’ and L5-6’ loops. In particular, the aromatic side chains of Trp712 and Phe715 dock into a shallow groove formed by residues at the C-terminal end of H5 and the following loop. This arrangement is thus topologically unrelated to the canonical dimerization mode, in which H10-11 helices from two monomers run parallel to each other, resulting in much higher interaction areas of ∼1,000 Å^2^. Noteworthy, the side chain of Tyr545’ engages also in important contacts at this protein-protein interface, docking on H9 from a neighboring monomer. This larger interface is strengthened by salt bridges between residues Arg690 and Asp549’ and by several H-bonds (e.g., between the carbonyl oxygen of Phe774 and the hydroxyl of Ser550’). A second, symmetric homodimer is centered on the aromatic side chains of Tyr638 (L6-7) and Phe735/Tyr738 (C-terminal end of H11) facing each other. However, since positions 638 and 738 are occupied by smaller polar residues in WT GR (Cys and Gln, respectively; Fig. 1C), this arrangement is unlikely to be significant *in vivo*.

Finally, two related, medium-resolution structures of GR-LBD·DEX bound to SHP were obtained in the tetragonal and cubic space groups (I4122 and I4132, respectively; Figs. 2C, D). Also in this case, a symmetric homodimer is observed in which the Tyr545/Tyr545’ aromatic rings are stacked, although the overall arrangement of LBD modules differs strongly from the Tyr545-directed dimer found in C2 crystals. Additional H1-H3’ contacts result in a more compact conformation, which is stabilized by H-bonds between both main- and side-chain atoms of the two monomers, including a Glu542-Arg569’ salt bridge. The largest interaction interface in these crystals features a trimeric arrangement in which loops L1-3’/H3’, S2-L6’ and L11-12’ dock perpendicularly onto H10-11. The large, buried surface area in this trimer appears to compensate the electrostatic repulsion of abutting Asp641 carboxylates from the three monomers around a pseudo (in the tetragonal form) or exact 3-fold axis (in the cubic cell).

A complete catalogue of homodimeric arrangements illustrates the multivalent potential of the GR-LBD. The fact that even minor changes in protein complexes and crystallization conditions result in different GR-LBD arrangements, as demonstrated by the variety of crystal contacts described above (Figs. 2A-D), prompted us to systematically analyze protein-protein contacts in all crystal structures of the domain previously deposited in the PDB. The results of this analysis are summarized in Figs. 3A, B and Supplementary Tables 2 and 3. GR-LBD residues involved in homodimer formation cluster in four areas on the protein surface, the “front”, “back”, “top” and “base” of the domain (Figs. 3B-D), in the standard view of NRs shown in the center of Fig. 1B.

**Fig. 3.**
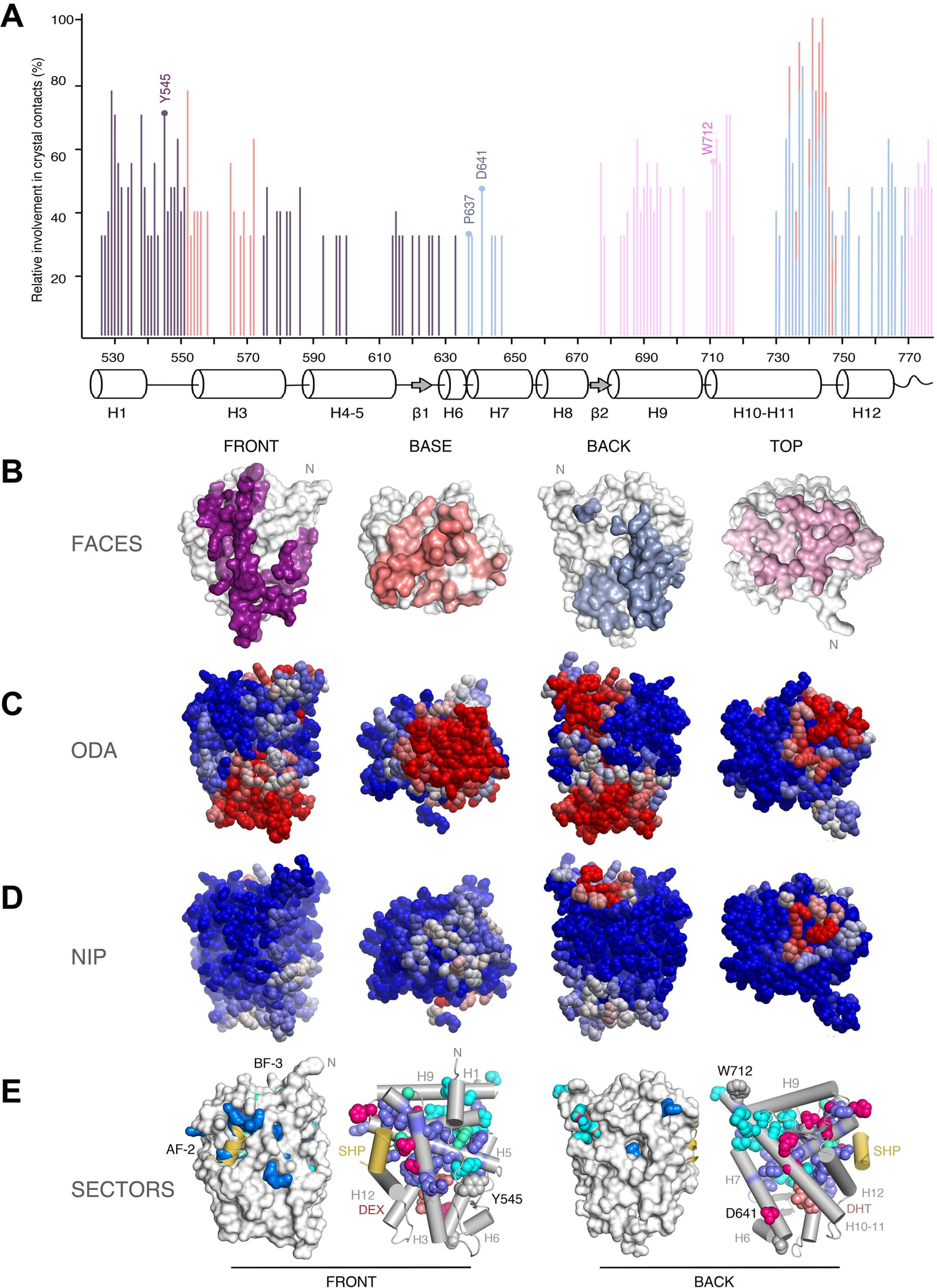
legend: Experimental structures and bioinformatics analyses unveil four major homodimerization surfaces on GR-LBD. **A**, Relative frequencies of residue involvement in GR-LBD homodimer formation. Bar height indicates how often a given residue engages in crystal contacts in all available structures of GR-LBD, normalized to the residue most frequently found in homodimer interfaces, Leu741. Secondary structure elements given below the plot correspond to the crystal structure of human GR-LBD resolved at the highest resolution, 6NWL. **B**, Residues involved in GR-LBD homodimerization cluster in continuous patches on its front (colored purple), base (coral), back (blue) and top (pink) faces. The association of these four faces yields the catalog of GR-LBD dimers represented in Fig. 4. Models are shown in the same orientation and at the same magnification in panels **C-E** below. **C**, Predicted protein-protein interaction optimal docking areas (ODA). ODA “hotspots” (residues with favorable docking energy; ODA < −10.0 kcal/mol) are colored red, residues with ODA > 0 kcal/mol are shown in blue, and intermediate values are scaled accordingly. ODA hotspots form continuous surface patches that essentially overlap with the four protein-protein interaction interfaces shown in panel **B. D**, Hotspot interface residues predicted from docking experiments. Surface residues are colored according to their normalized interface propensities (NIP). Residues with NIP > 0.4 and < 0 are colored red and blue, respectively; intermediate values are scaled accordingly. **E**, Statistical coupling analysis (SCA) identifies two sectors of clustered, physically connected residues in GR-LBD. The front and back orientations of GR-LBD are depicted, and in both cases the module is represented as a solid surface and as a cartoon, with helices shown as rods and labeled. Residues belonging to sectors I and II are shown with their side chain atoms as spheres, colored cyan and dark blue, respectively. Other important residues are also shown for orientation and labeled. The AF-2-bound SHP peptide is colored yellow, and the DEX ligand is represented as salmon spheres.

Next, we analyzed which combinations of these four homodimerization interfaces have been encountered in crystal structures. This analysis revealed 20 topologically distinct homodimers, numbered #1-20 throughout the manuscript (see Supplementary Table 3 for interacting residues in all monomer pairs). These homodimeric arrangements appear to cover the whole GR-LBD self-association landscape. Along with 11 symmetric (isologous) dimeric arrangements (i.e., between the same secondary structure elements / residues, such as in the Tyr545-mediated dimers described above), asymmetric or heterotypic homodimers (i.e., where the contacting GR-LBDs engage in interactions using different elements) are also common (9 arrangements).

We further explored the homodimerization potential of GR-LBD with a state-of-the-art protein-protein docking procedure. A total of 12,000 docking dimers were generated using the coordinates from PDB entry 5UFS. This analysis revealed the existence of at least one docking orientation close to 16 of the 20 representatives “crystal homodimers” (with a root-mean-square deviation (RMSD) ≤ 10 Å). Interestingly, the 2nd best-scoring docking orientation was close (7.3 Å RMSD) to one of the dimers (#20). In two further cases (#8, #10), there were docking orientations within 5 Å from the crystal structures, although with no optimal docking scoring.

For an unbiased estimate of the similarity between different homodimeric conformations we first considered the overlapping of shared contact residues. To this end, we mapped these sets of residues into a multiple sequence alignment and calculated distances between the resulting vectors using different metrics (see Fig. 4A for the clustering obtained using Jaccard’s similarity index). This analysis confirmed e.g., the topological similarity between two front-to-front homodimers: the first described non-canonical conformation (#1, PDB 1M2Z (Bledsoe et al, 2002)) and #2, an arrangement observed in PDB 4P6W (He *et al*, 2014) (Supplementary Tables 2 and 3). On the other hand, front-to-front homodimer pairs #6 and #11, although sharing interface residues, differ strongly in that the two monomers are arranged parallel and antiparallel to each other, respectively. Similarly, homodimer pairs #9 and #10 share the important residue Ile628 at the center of their intermonomer interfaces, but the two modules are quite differently oriented relative to each other.

**Fig. 4.**
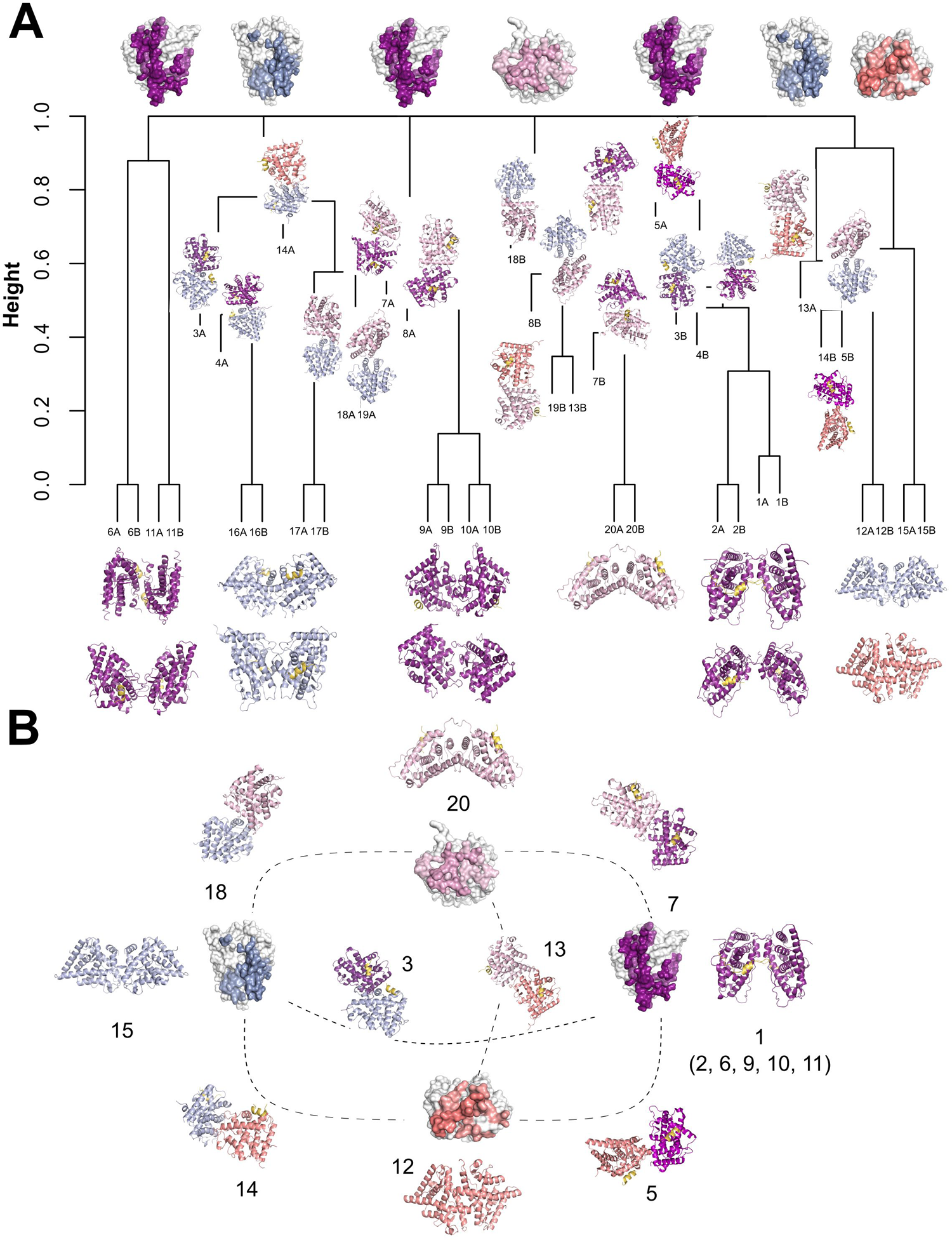
legend: An integrated catalog of GR-LBD homodimers. The four distinct GR-LBD protein-protein interfaces associate to generate 20 topologically different homodimers. **A**, A dendrogram based on a hyerarchical analysis of protein-protein contacts using Jaccard’s index groups the 20 unqiue GR-LBD asemblies into six different clusters. **B**, Relationsships between the different GR-LBD homodimers. For orientation, monomers highlighting the four interacting surfaces are placed at the cardinal points in this panel (top, nord; front, east; base, south; and back, west), colored-coded as in Fig. 3. Monomers in 10 representative homodimers are depicted as cartoons; each monomer is colored according to the face used to associate with its partner. Dimers are placed closest to the generating monomers.

Alternatively, GR-LBD homodimers were superimposed on a common origin and classified by mapping the centers of coordinates of their interaction surfaces (Supplementary Fig. 3A). Supplementary Fig. 3B shows the orientations of these surfaces, represented by vectors between the common center of coordinates and each interaction surface. A hierarchical clustering analysis based on the Euclidean distance between these vectors grouped all homodimers into six clusters (Supplementary Fig. 3C). From the spherical coordinates of the vectors representing the orientations of the interaction surfaces (sinusoidal equal-area projection in Supplementary Fig. 3E), we found that these clusters can be associated with combinations of the previously defined surfaces: cluster 1 (top-front), cluster 2 (top-back), cluster 3 (base-back), cluster 4 (base-front), cluster 5 (front), and cluster 6 (base). While each cluster may contain surfaces with different binding energy values (Supplementary Fig. 3D), arrangements corresponding to top and back interaction surfaces have in general more favorable binding energy. The distribution of interfaces in the top 1,000 docking models also shows significant clustering around regions with favorable binding energy (Supplementary Fig. 3F). However, docking solutions also clustered in other regions with less optimal energy, such as front and base surfaces.

### Two sectors define the internal circuits linking major interaction sites in the GR-LBD

To identify residues responsible for the functional specificity of GR we first run multiple correspondence analysis (MCA), which did not replicate the previously reported family classification (Weikum *et al*, 2017b), and only identified three residues that divide the NR superfamily into two clusters. This prompted us to use more sophisticated statistical tools to search for evolutionary conserved units in GR-LBD. To analyze if self-association surfaces may be allosterically coupled to other functional regions, we performed a statistical coupling analysis (SCA), which entirely relies on correlated amino acid variations across the domain without considering its 3D structure (Halabi *et al*, 2009; Lockless & Ranganathan, 1999). Indeed, this analysis identified 40 residues that decompose the GR-LBD sequence into two quasi-independent groups of correlated residues or “sectors” (Fig. 3E).

Sector 1 comprises 17 residues in and around H1 and H10, most notably LBP residues Met601 and Arg611 along with the nearby Phe606, whereas sector 2 features 20 residues mostly from H3 and H5 (e.g., Met604 in the LBP, Lys579, Phe584, Gln597 of AF-2, and Trp577 in an internal path connecting AF-2 to the LBP). Finally, three residues (Gly583 of BF-3, Leu596 at the floor of the AF-2 groove, and the internal Tyr663) belong to both sectors. Interestingly, all these residues are clustered in the upper half of the domain, where both sectors are physically interconnected (Fig. 3E). Sector 1 residues cluster around the N-terminus of the domain and are thus likely candidates to interact with hinge residues and the preceding DBD. Perhaps more relevantly, sector 2 comprising residues profusely innervate the LBP and AF-2 regions while Arg611 from sector 1 is essential to position hormones in the LBP. The three residues that belong to both sectors are strategically located to cross-connect the LBP with AF-2 and BF-3 pockets. Taken together, our results suggest that both sectors link functionally relevant regions thus coupling e.g., ligand binding to coregulator binding or chaperone docking/release.

### *In vitro* crosslinking experiments corroborate non-canonical dimerization of GR-LBD

The results presented above suggest that many surface-exposed residues of GR-LBD engage in a variety of crystal contacts. To clarify which structural elements/residues might be involved in homodimer formation in solution, we took advantage of the observation that some of the crystal interfaces are stabilized by intermolecular H-bonds between Glu/Asp carboxylates and Lys ammonium groups, or that such bonds could be easily formed upon side-chain rotations, and that these linkages can be “frozen” upon incubation with the zero-length crosslinker, EDC.

To verify whether some of these Asp/Glu-Lys H-bonds are formed in solution, we incubated GR-LBD in the presence of EDC. Indeed, we observed rapid formation of a covalent dimer, as well as a fainter band corresponding to a tetrameric arrangement(s) (Fig. 5A). To identify charged residues responsible for EDC crosslinking, bands corresponding to oligomeric GR-LBD forms were excised from the gels, subjected to enzymatic digestions with either trypsin or chymotrypsin, and analyzed by mass spectrometry (MS) (Supplementary Fig. 3A and Supplementary Tables 4 and 5). Most notably, we found that elements essential for top-to-top (#20) and front-to-front (#1, #2, #6, #9, #10 and #11), non-canonical dimerization are overrepresented among EDC-linked peptides, with the most common contacts involving (1) residues of H1 and L1-3, on the one side, and from H3, on the other, which would correspond to front-to-front interactions, as well as (2) H9 and the L9-10 linker from two monomers, which is compatible with homodimer #20 (Fig. 5B, Supplementary Fig. 3B and Supplementary Table 4). Similar results were obtained with the MS-cleavable, urea-based crosslinker, DSBU (Supplementary Table 6).

**Fig. 5.**
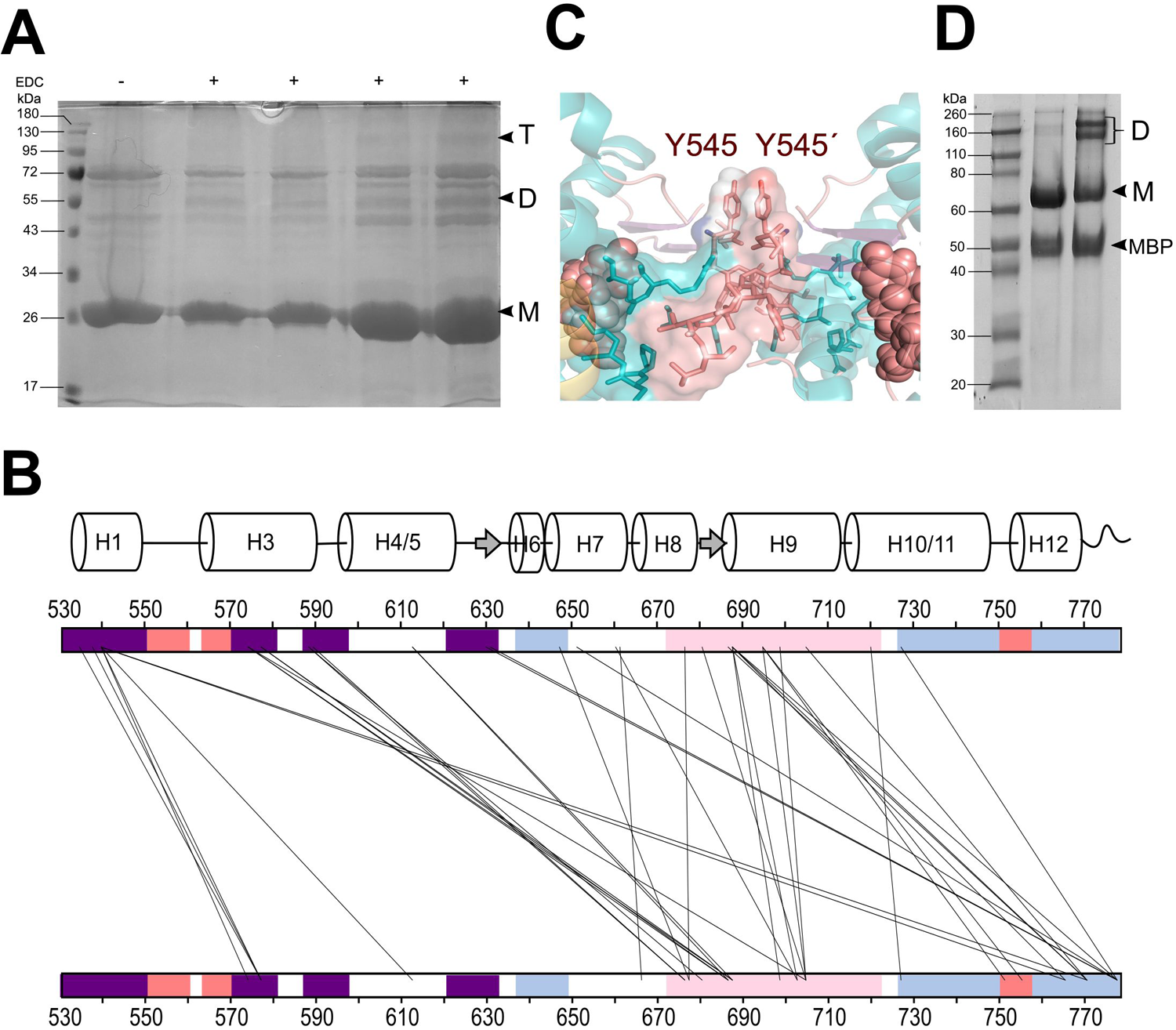
legend: Several GR-LBD homodimers are populated in solution. **A**, SDS-PAGE analysis of GR-LBD samples after incubation with the zero-length crosslinker, EDC. Notice bands with relative molecular masses corresponding to GR-LBD dimers (D) and tetramers (T) in all the lanes except control lane 1 (no EDC added). Lanes 2 and 3, protein incubated at about 0.37 mg/ml; lanes 4 and 5, protein incubated at about 1.5 mg/ml. Samples in lanes 2 and 4 were treated at room temperature; those in lanes 3 and 5 at 30 °C. **B**, Crosslink map of EDC-treated GR-LBD showing all crosslinked peptides captured. Regions corresponding to the top, front, base, and back surfaces are colored as in Figs. 3 and 4. A secondary structure plot is shown above the map. **C**, Closeup of the major homodimerization interface in C2 crystals (front-to-front homodimer #11), dominated by stacked phenol rings of Tyr545/Tyr545’ residues. **D**, Non-reducing SDS-PAGE analysis of purified GR-LBD(Y545C) (lane 3) shows spontaneous dimerization in solution. Note that the WT protein does not form dimers when incubated at the same concentration (lane 2).

### Cysteine point mutant Y545C demonstrates non-canonical homodimerization of GR-LBD in solution

Inspection of homodimer interfaces in GR-LBD revealed several symmetric arrangements in which the side chain of a solvent-exposed residue from one monomer is located within VdW distance of the same residue from a crystal neighbor (Figs. 2A and 5C). To verify whether some of these conformations are populated in solution, we have generated several cysteine point mutants of the GR-LBD. All studied mutants were properly folded, as indicated by only minor decreases in melting temperatures in differential scanning fluorimetry (DSF) analysis (not shown), in line with the results of a systematic bioinformatics analysis of mutant stability performed using Fold-X (not shown). Incubation of purified GR-LBD(Y545C) in low-reducing conditions resulted in the rapid formation of covalent dimers (Fig. 5D). By contrast, neither the WT protein nor other Cys mutants tested (e.g., D641C, S744C) dimerized under the same conditions. These findings strongly suggest that the side chains of Tyr545 from two monomers are close enough in solution, at least in a subset of GR-LBD molecules.

To directly proof that residues Cys545/Cys545’ are responsible for disulfide bridge-mediated dimerization in solution, bands corresponding to the dimer were excised from the gel, treated with iodoacetamide to block free Cys residues, and subjected to enzymatic digestion with trypsin and GluC. MS analysis of these digests allowed indeed the identification of peaks corresponding to peptide V^543^LCSGYD^549^ crosslinked to either V^543^’LCSGYD^549’^ or V^543’^LCSGYDSTLPDTSTR^558^’, thus confirming Cys545-mediated covalent bond formation (Supplementary Fig. 3C; see also Supplementary Table 7 for a list of a, b and y ions that allowed unambiguous identification of the crosslinked peptides).

To further assess the contribution of Tyr545 to GR homodimerization in solution, SPR assays essentially equivalent to those described above for WT GR-LBD were performed with its Y545C and Y545A mutants. Indeed, experiments conducted with the Cys mutant revealed significant increases in affinity. The increase was highest when the mutant was used as both ligand (i.e., chip-immobilized) and analyte, and affects mainly the first binding site, with a 3-fold increase in affinity (kD1 = 0.8 vs. 2.2 µM for the WT-WT association; Supplementary Fig. 2B). By contrast, presence of a less bulky alanine at position 545 led to a slightly less tight association (Supplementary Fig. 2C). We also crystallized and solved the structure of the GR-LBD(Y545A) variant, thus confirming proper folding of the generated point mutants. Noteworthy, Y545A crystallized in the P61 space group, which does not involve symmetric contacts between the side chains of residues at position 545 (Fig. 2B). Altogether, our results confirm that residue Tyr545 plays an important role in GR-LBD homodimerization in solution.

### Residues Tyr545 and Asp641 modulate multimerization of full-length GR

Quantitative fluorescence microscopy in living cells (the number and brightness method, N&B) allows to estimate the average oligomeric state of a fluorescent protein from its molecular brightness (ε; Digman et al, 2008). For N&B experiments we routinely use GR^null^ mouse adenocarcinoma cells, which possess a tandem array of DNA binding sites for GR, the MMTV array. Cells are transfected with GFP-labeled mouse GR (GFP-mGR) or variants thereof, and the oligomeric state of fluorescently tagged GR molecules is quantified by comparing to a constitutively monomeric GR variant (N525*). Further, presence of the MMTV array allows us to differentially assess oligomerization of the nuclear receptor in the entire nucleoplasm and in a region highly enriched in specific binding sites.

To prove the relevance for the full-length receptor of key surface-exposed residues identified in GR-LBD homodimer interfaces in vitro, we generated alanine mutants of mice GR at positions topologically equivalent to human residues Tyr545, Pro637, Asp641 and Trp712, all of which are conserved in ancGR2 (Fig. 1C). Further, all residues but Asp641 are strictly conserved from fish to humans, and Asp641 is conservatively replaced by a glutamate in non-mammals (Supplementary Fig. 1). To study the impact of the Tyr545→Ala exchange in the background of two other variants previously shown to be important for GR homodimerization (Presman *et al*, 2016), we also generated the double mutant (Tyr545Ala, Ile628Ala) (in following termed GR^dim/Y545A^) and the triple mutant (Ala458Thr, Tyr545Ala, Ile628Ala), or GR^mon/Y545A^. Finally, we also generated double mutants in which a GR variant that tetramerizes both in the nucleus and at the array (Pro474Arg, Paakinaho et al, 2019) was combined with either P637A or D641V. (These variants, (Pro474Arg, Pro637Ala) and (Pro474Arg, Asp641Val), are in following termed GR^tetra/P637A^ and GR^tetra/D641V^, respectively. All mutated residues in the DBD or LBD moieties are highlighted in Fig. 6A.

**Fig. 6.**
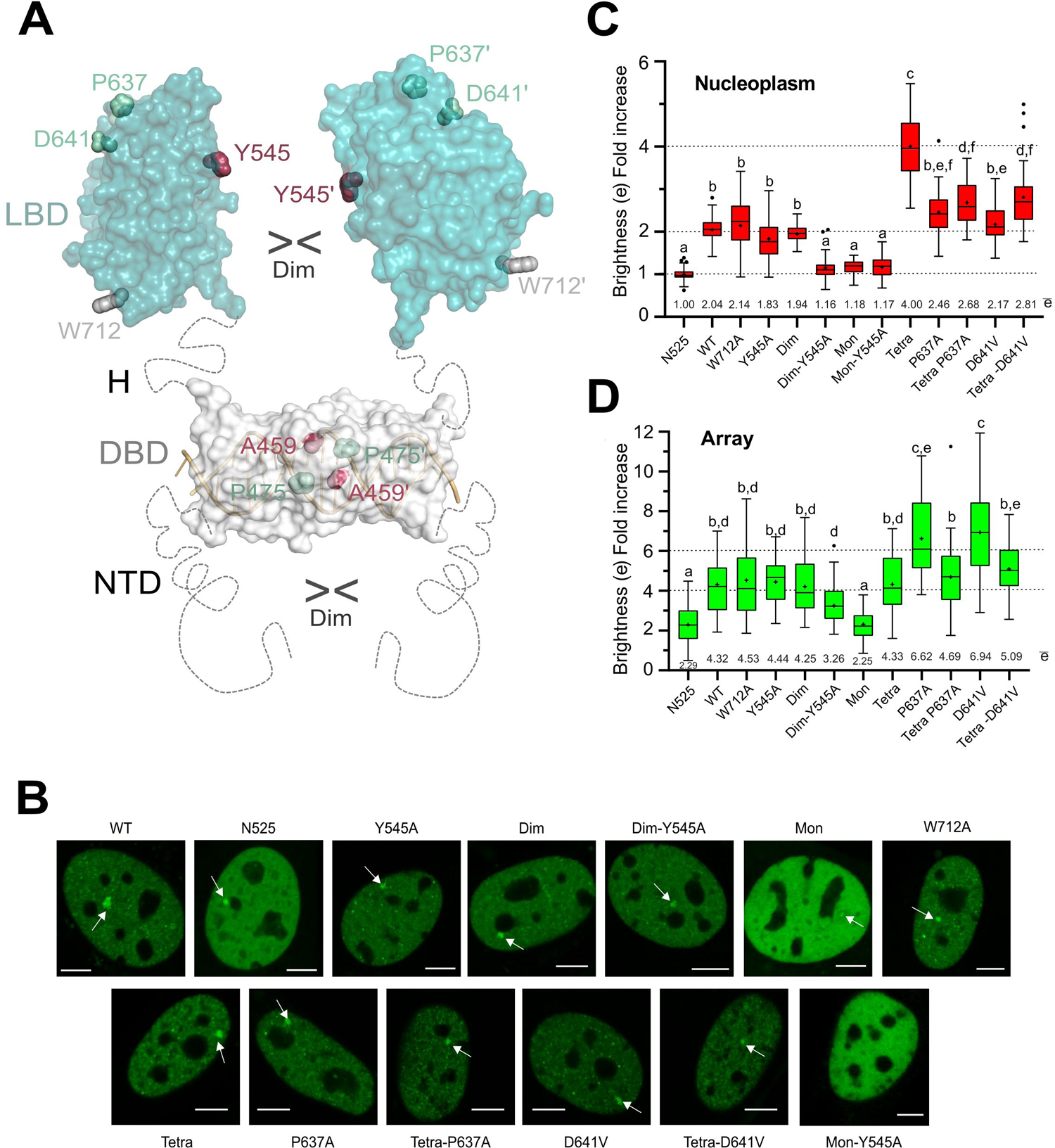
legend: Mutation of LBD-LBD interface residues profoundly affects the multimerization behavior of full-length GR. **A**, Schematic 3D model of the full-length protein. The intrinsically disordered NTD mediates liquid-liquid phase separation (LLPS) and is followed by the globular DBD (here, the structure of its DNA-bound dimer is shown), and by the actual LBD. Note that the length and flexibility of the DBD-LBD linker (hinge, H) allows for the formation of various different homodimers. The side chains of all residues mutated to assess the multimerization behavior of GR are shown as spheres. **B**, Subcellular localization of WT GFP-mGR and indicated mutants in 3617-GRKO cells, as assessed by fluorescence microscopy. Variant N525* lacks the entire LBD and remains monomeric (Presman *et al*, 2016). White arrowheads point to the MMTV arrays. Scale bar: 5 µm. Data for WT_GR, GR^dim^, GR^mon^ and GR^tetra^ were taken from (Presman *et al*, 2016) and are shown for comparison purposes. Residue numbers correspond to the human protein to facilitate comparisons. **C**, GR quaternary structure in the nucleus, as determined in N&B assays. The fold increase in molecular brightness (ε) relative to the N525* monomeric control is shown. **D**, Quaternary structure of DNA-bound GR. The results of N&B assays at the MMTV arrays are represented as in panel C. Note that simultaneous disruption of intermonomer interactions mediated by the DBD (Ala458Thr) and the LBD (Tyr545Ala) in the GR^dim/Y545A^ double mutant results in a variant that is monomeric in the nucleus, while at the array it formed mostly trimers. In panels **C** and **D**, centered lines show the medians and crosses represent sample means (average numbers below each box-plot). Box limits indicate the 25th and 75th percentiles; whiskers extend 1.5-fold the interquartile range from the 25th and 75th percentiles, with outliers represented by dots. Boxes with different superscript letters are significantly different from each other (p<0.05; one-way ANOVA followed by Tukey’s multiple comparison test).

Next, we transiently transfected GFP-mGR and the generated mutants into GR^null^ mouse adenocarcinoma cells and performed N&B experiments as previously described (Presman *et al*, 2017, 2016; Digman *et al*, 2008). All mutants translocate to the nucleus upon hormone stimulation (Fig. 6B), indicating proper folding and unaffected ligand binding. Further, all variants except those carrying the GR^mon^ double mutant (Ala458Thr, Ile628Ala) were visible at the MMTV array, suggesting that DNA binding was not impaired either (Fig. 6B, arrowheads). Severely reduced genome-wide chromatin binding for GR^mon^ has recently been shown (Johnson *et al*, 2021), and only a very small percentage of GR^mon^ cells form visible arrays (Presman *et al*, 2016, 2014). Interestingly, we did not detect any arrays in experiments with GR^mon/Y545A^ (Fig. 6B), suggesting an even more drastic phenotype for this triple mutant.

While W712A oligomerizes as the WT receptor both in the nucleoplasm and at the array (Figs. 6C, D), Tyr545Ala substitution appears to slightly decrease dimerization in the nucleoplasm (ε= 1.83). By contrast, Pro637Ala produces a slight increase in oligomerization (ε= 2.46), even though neither difference achieves statistical significance. Since FL GR dimerizes at least through both DBD and LBD moieties (Fig. 6A), we tested the effect of the Tyr545Ala mutation in the GR^dim^ background, which has impaired DBD-DBD contacts yet mostly dimerizes in the nucleoplasm (Fig. 6C, Presman *et al*, 2014). Indeed, the GR^dim/Y545A^ double mutant shows significant tendency to remain monomeric in the nucleus (ε= 1.16), like the previously characterized GR^mon^ (Fig. 6C, Presman *et al,* 2014). Taken together, these results confirm the important role of Tyr545 in the dimerization of GR in live cells.

On the other hand, the Asp641Val mutation, linked to Chrousos syndrome (Hurley *et al*, 1991) and the Pro637Ala substitution promoted higher-order oligomerization at the array (Fig. 6D), possibly hexamers or a mixture of tetra- and octamers. These findings prompted us to analyze the impact of these two variants when combined with a DBD mutation that enforces GR tetramerization, Pro474Arg. Unexpectedly, instead of synergizing both mutants reversed GR^tetra^ oligomerization in the nucleus (Fig. 6C). By contrast, the GR^tetra^ mutation abrogated the ability of P637A and D641V variants to form higher-order oligomers at the array level (Fig. 6D). These observations highlight a complex relationship between the different structural domains of the NR.

## Discussion

Although many structures of GR-LBD have been reported in a wide variety of crystal forms (Supplementary Table 2), the physiologically relevant conformation(s) of GR and other oxosteroid NRs remain disputed. The new structures presented here highlight the ability of different GR-LBD surfaces to engage in homophilic contacts, resulting in different quaternary arrangements depending on the bound agonists/antagonists, cofactors, and other biochemical parameters. For instance, a monoclinic structure of ancGR2-LBD bound to another synthetic GC, triamcinolone acetonide, and complexed to a shorter SHP peptide had been previously reported (PDB 5UFS; Weikum et al, 2017b). Interestingly, 5UFS features a Tyr545-centered parallel dimer almost identical to the topologically equivalent arrangement in our current C2 crystals (Fig. 2A and Fig. 5C). In the case of the hexagonal crystals, similar structures of ancGR2-LBD bound to either DEX or a different GC (mometasone furoate) and complexed to a TIF-2 peptide had been previously reported (PDB entries 3GN8 and 4E2J, respectively). Although there are only relatively small differences in the cell constants compared to the current P6_1_ structure (a and b axes are ∼5% longer in our crystals, while the c axis is ∼4% shorter), this results in a markedly different small intermonomer interface, which is asymmetric in 3GN8/4E2J.

These and other observations prompted us to dissect the oligomerization capability of GR-LBD. Careful inspection of intermonomer contacts in all available 3D structures allowed us to identify 20 topologically different homodimeric architectures (Fig. 4 and Supplementary Table 3). These experimental GR conformations could be grouped into six different clusters considering relationships between interacting residues (Fig. 4A), which correspond to three partially overlapping though topologically distinct front-to-front homodimers (#1, #2, #6, #9, #10 and #11), along with base-to-base (#12), top-to-top (#20), and back-to-back (#15, #16 and #17) arrangements. A similar pattern emerges when these homodimers are represented in spherical coordinates (Supplementary Fig. 4E) or using an unbiased graph analysis (Supplementary Fig. 4C).

Next, we analyzed the behavior of WT GR-LBD and several point mutants generated according to structural and functional evidence in solution, coupled with quantitative fluorescence microscopy of FL GR and several mutants in cells. The results of these investigations, under careful consideration of geometric and energetic parameters, as well as conservation of major interface residues, allow us to postulate the more likely quaternary arrangements of GR-LBD modules associated with its different pathophysiological roles.

### GR does not dimerize in an AR-like conformation

It has long been accepted that the “canonical”, H10-mediated homodimeric conformation adopted by ERα-LBD (Supplementary Fig. 5B) is not possible in oxosteroid receptors, due to both non-conservative replacements of essential interface residues and partial occlusion of the dimerization interface by the F-domain3–5,13,50. Several observations suggested that the non-canonical dimerization mode we have recently reported for AR (Supplementary Fig. 5A), centered on H5 and neighboring elements instead (Nadal *et al*, 2017; Jiménez-Panizo *et al*, 2019; Fuentes-Prior *et al*, 2019) could be adopted by other members of the subfamily. For instance, aromatic residues involved in maintaining the rigid, dimerization-competent structure of H5 such as Trp610 and Tyr613 (in human GR) are highly conserved in all oxosteroid receptors.

However, in contrast to expectations none of the 20 GR-LBD homodimers can be considered as topologically equivalent to the one observed in the crystal structure of AR-LBD, illustrating a more complex multimerization behavior than previously anticipated. Replacement of AR interface residues such as Thr656 by positively charged Lys/Arg residues and of the following Asn657 by bulkier Gln/His residues in all other members of the subfamily might preclude formation of AR-like homodimers in GR / MR / PR. This highlights the difficulty to extrapolate the quaternary structure of a given NR to other, even closely related family members, and the need for experimental evidence to identify physiologically relevant conformations (Fuentes-Prior *et al*, 2019). A related arrangement had been previously observed in some structures of agonist- or antagonist-bound GR-LBD (Bledsoe *et al,* 2002), but its biological relevance has been repeatedly questioned (Kauppi *et al*, 2003), and several unrelated multimeric assemblies could be postulated (discussed in detail below).

Possible quaternary arrangements of the GR-LBD and their functional relevance *in vivo*.

The largest homodimeric interfaces correspond to antagonistic conformations of GR. The symmetric back-to-back (#15) and base-to-base homodimers (#12) have much larger interface areas (∼1470 and 1200 Å^2^, respectively) and much lower energies (−98.0 / −83.0 kcal/mol) than all other GR-LBD conformations (750 Å^2^ and −50.0 kcal/mol for the next best configuration; Supplementary Fig. 4D). Interestingly, these dimers are only observed in GR-LBD bound to the antagonist, mifepristone/RU-486 (PDB entries 1NHZ and 5UC3, respectively, Kauppi *et al*, 2003), and share several important features. First, the bound antagonist enforces displacement of the C-terminal H12, which partially disrupts the LBP. More importantly, in both cases the AF-2 pocket of one monomer is partially covered by either H12 (#15) or H3 (#12) from a neighboring molecule, thus interfering with coregulator binding (Fig. 7A, B). Further, essentially the same arrangements are found in the crystal structure of the dominant negative GRβ, which is known to bind only antagonists53.

**Fig. 7:**
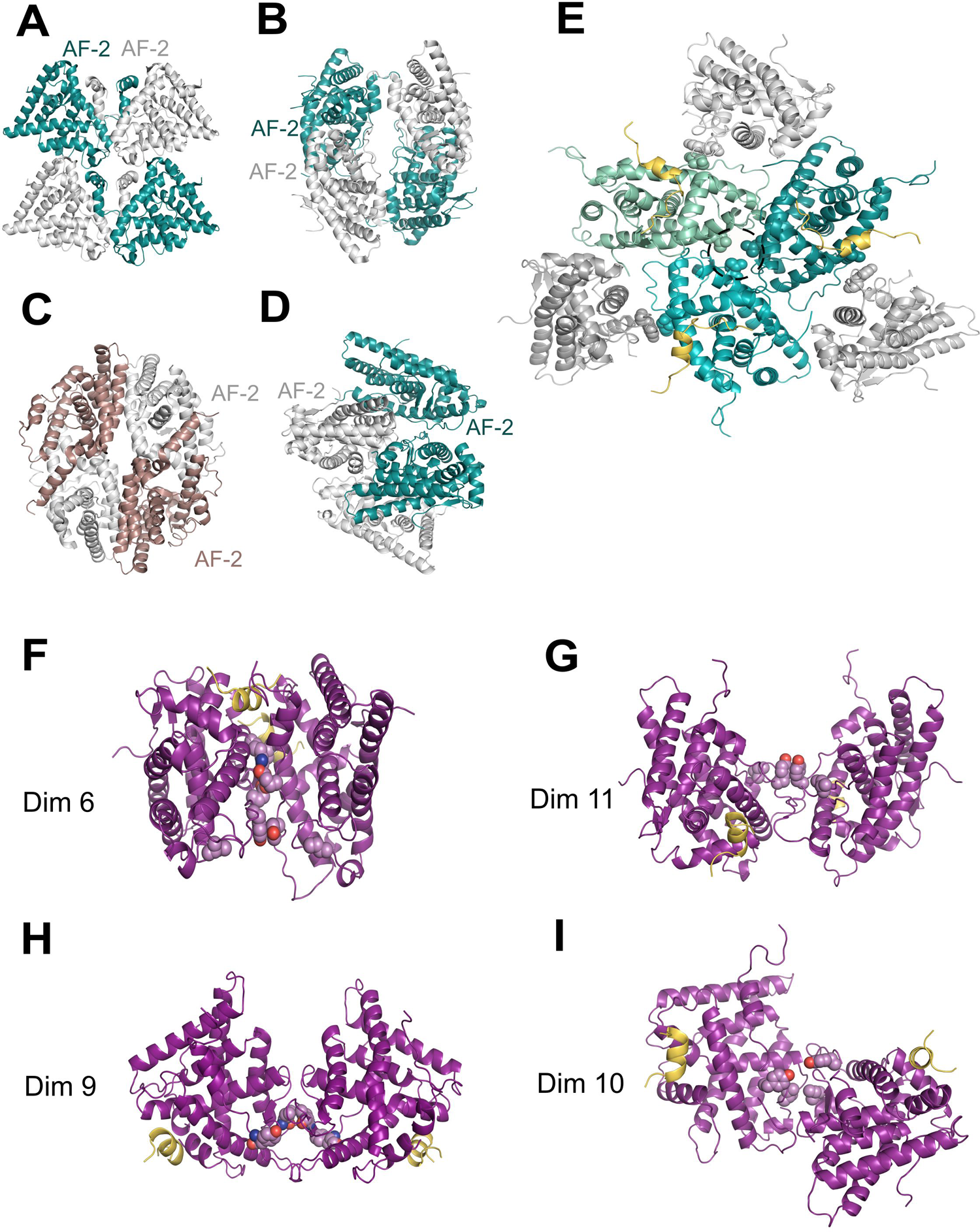
Multimeric arrangements of GR-LBD compatible with current structural information. **A-D**, Tetrameric arrangements found in the crystal structures of GR-LBD bound to the potent antagonist, RU-486 (PDB codes 1NHZ (**A**) and 5UC3 (**B**), respectively) compared to (**c**) the inactive RXR-LBD tetramer (1G1U). Two-fold axes generating the dimers-of-dimers run roughly perpendicular to the page plane. **D**, Model of GR-LBD tetramer generated by docking dimer #11 onto itself. The model is compatible with the results obtained with the Y545C mutant and EDC-crosslinking assays. **E**, Putative GR-LBD hexamer favored by the Asp641Val mutation. The central trimer corresponds to an arrangement observed in tetragonal and cubic crystal forms (#14; Figs. 2c, d and 4), while the external monomers dock according to conformation #4. Asp641 residues from the central trimer are encircled. **F-G**, Models of human GR-LBD homodimers based on the observed assemblies #10 (**F**), #6 (**G**), #9 (**H**), and #11 (**J**). The critical homodimerization residues, Tyr545 and Ile628, are shown as color-coded spheres in all cases.

Interestingly, RU-486–bound GR-LBD forms a covalent homodimer through disulfide bridge Cys736/Cys736’ (in conformation #15; Kauppi *et al,* 2003). The fact that we have not observed this covalent dimer of WT GR-LBD in the presence of dexamethasone suggests that this quaternary arrangement is exclusive of the antagonist-bound receptor. Further, tetrameric arrangements in 1NHZ/5UC3 and in a third structure of mifepristone-bound GR (3H52; Schoch *et al*, 2010) are not compatible with the results of our XL experiments with EDC (Fig. 5A and Supplementary Fig. 2B), or engage the Trp712 side chain, which has no effect on GR homodimerization (Fig. 6C, D), or involve important partial unfolding events that result in swapping of N- and/or C-terminal residues between two monomers (up to L1-3 or from H12 on, respectively). Of note, similar rearrangements have not been observed in the crystal structures of core NRs bound to agonists and their DNA response elements (Chandra *et al*, 2017; Fuentes-Prior *et al*, 2019). Altogether, these findings strongly suggest that the most stable, back-to-back (#15) and base-to-base conformations (#12) of GR-LBD are associated with inactive, (self-repressed receptor states, which are induced or stabilized by antagonist binding to the LBP. Preferential binding of corepressor peptides to these antagonist-bound conformations has been reported (Schoch *et al*, 2010; Min *et al*, 2018), but the significance of these findings in the context of full-length cofactors is not clear. Noteworthy, inactive RXRα adopts a disc-like tetrameric conformation, in which H12 from one molecule protrudes away and docks on the AF-2 pocket of an adjacent monomer (Fig. 7C; Gampe *et al,* 2000). Although the overall arrangement differs from the ones adopted by GR-LBD, the fact that this important surface pocket is occluded in both cases points to a general feature of self-repressed or inactive conformations in NRs. Interestingly, also in the inactive RXR tetramer two monomers are covalently linked through a disulfide bond, suggesting yet unexplored connections between redox state and NR biology.

### Formation of non-physiological multimers appears to underlie the deleterious effect of Chrousos syndrome mutants

Unexpectedly, in our N&B experiments we observed that a GR mutant previously linked to Chrousos syndrome, p.Asp641Val49, preferentially formed higher-order oligomers when bound to DNA (Fig. 6D). This non-conservative mutation has a particularly strong stabilizing impact of 3.4 kcal/mol for each dimer pair on the GR-LBD trimer observed in our tetragonal and cubic crystals (#14; see Fig. 2D and Supplementary Table 3). The positive effect of the mutation results from the relief of strong electrostatic repulsion between the three abutting Asp641 residues, coupled with favorable VdW contacts made by the aliphatic Val641 side chains with each other and with Cys638 from a neighboring molecule. Notably, conformation #4 is fully compatible with this trimer and generates a closed hexamer by additional interactions between the N-terminal end of H10 and H12’’ at a third interface (Fig. 7E). This conformation would occlude the AF-2 pockets of the “external” monomers and is thus incompatible with coregulator binding.

Interestingly, several additional residues associated with Chrousos disease are located at or close to one of the three monomer-monomer interfaces, and their mutations might promote local conformations that also stabilize this hexamer. For instance, replacement of Thr556 by an aliphatic isoleucine would stabilize this arrangement through contacts with e.g., residues Met560 and Pro637 from its “own” monomer and/or His645’/Asn731’ from a neighboring molecule. Similar considerations apply to mutations such as Arg714Gln, Phe737Leu, Ile747Met and Leu773Pro as well as to the variant of the central interface Pro637Ala, which also forms higher-order oligomers at the array (Fig. 6D). Thus, we conclude that not only p.Asp641Val but also other GR variants associated with Chrousos disease stabilize multimeric forms that are incompatible with active GR tetramers on DNA. This is, to the best of our knowledge, the first time in which non-productive multimers of a NR are postulated as the molecular basis of a human disease.

A three-fold reduced affinity of the Val641 mutant for DEX was previously proposed as the underlying molecular defect in the D641V variant (Hurley *et al*, 1991). However, since Asp641 is exposed on the surface of the protein and is located 8 Å away from the closest DEX atom, it is unlikely to play any relevant role in hormone recognition. Accordingly, nuclear translocation of FL GR(D641V) in live cells was comparable to WT GR, suggesting no major ligand affinity issues. Residue Asp641 was not predicted as part of a sector, which seems to exclude also indirect (allosteric) effects transmitted to the LBP.

### The N-terminal end of H10 and surrounding residues are important for chaperone-binding to GR but are unlikely to play a major role in receptor multimerization

Because of their ability to engage in important protein-protein interactions, the bulky, aromatic Tyr/Trp residues are quite common at both homo- and heterodimeric interfaces57. GR-LBD features several exposed Tyr/Trp residues, two of which, Tyr545 and Trp712, are repeatedly found at monomer-monomer interfaces (Fig. 2 and Supplementary Table 3) and are strictly conserved from fish to humans (Supplementary Fig. 1). To explore the possible role of these residues in GR multimerization *in vivo*, we generated the Y545A and W712A mutants of FL GR and studied their behavior using quantitative fluorescence microscopy. Against expectations, elimination of the bulky indole ring at position 712 had no effect on the oligomerization properties of GR, and the W712A variant behaved as WT both in the nucleoplasm and DNA-bound (Fig. 6C, D). These findings strongly suggest that Trp712 is not important for GR multimerization *in vivo*. This would exclude, among others, the symmetric homodimer in which the Trp712 indole rings of two monomers occupy a shallow pocket between H9 and the F domain from a neighboring molecule (homodimer #20), and which had been proposed as the most likely conformation of the GR-LBD homodimer (Bianchetti *et al*, 2018).

Nevertheless, the fact that Trp712 participates in six topologically different homodimeric arrangements suggests an enhanced propensity for protein-protein interactions. Indeed, the recently reported cryo-EM structure of GR-LBD bound to the “client-maturation complex” Hsp90-p23 has revealed that Trp712 occupies the central position in a major binding epitope for the p23 co-chaperone (Noddings *et al,* 2020). However, none of the six GR-LBD homodimers that engage Trp712 appear to mimic the p23-GR heterocomplex, and there are no FXXMMXXM sequences in GR-LBD that would correspond to the GR-interacting helix in p23.

### Non-canonical GR homodimers centered on Tyr545 are critical for GR homodimerization

In contrast with the normal multimerization behavior of the W712A variant, alanine replacement of Tyr545 reduced receptor dimerization in the nucleoplasm. Moreover, when combined with an exchange that disrupts DBD-DBD contacts (Ala458Thr, GR^dim^), it resulted in a mostly monomeric form (Fig. 6C). Further, an important fraction of GR-LBD Y545C molecules rapidly and spontaneously forms covalent homodimers *in vitro* mediated by the Cys545-Cys545’ disulfide bond (Fig. 5C, D). Finally, alanine replacement of Try545 led to a reduced monomer-monomer affinity in SPR experiments (Supplementary Fig. 2E). Altogether these observations strongly suggest that the Tyr545 phenol ring plays a critically important role in GR homodimerization.

Two of the five topologically different homodimeric arrangements that involve the Tyr545 side chain (#7 and #8) are asymmetric and include the Trp712 indole ring; they are thus unlikely to be relevant *in vivo*. On the other hand, formation of the Cys545-Cys545’ disulfide bridge in the Y545C variant is compatible with two topologically different, roughly parallel and antiparallel symmetric homodimers (#6 and #11; Figs. 7F, G, respectively). Incomplete dimerization of GR-LBD(Y545C) suggests that other dimeric states are also populated in solution. Indeed, in a third, also antiparallel GR-LBD homodimer the Tyr545 side chains are located at the borders of the protein-protein interface (#9, Fig. 7H). Interestingly, this arrangement is centered on opposing residues Ile628/Ile628’, which are important for receptor dimerization (Presman *et al*, 2014, 2016). Finally, a second Ile628-centered, parallel conformation is found in crystals of ancGR1 (homodimer #10; Fig. 7I; Carroll *et al*, 2011). Thus, both parallel and antiparallel arrangements of LBD modules with different involvement of the Tyr545 and Ile628 “valences” are compatible with current experimental evidence and are equally possible in principle, as the associated buried surface areas and energies are similar (between 530 and 740 Å^2^ and between −8 and −26 kcal/mol). Some of these quaternary arrangements in solution could be verified in our XL-MS experiments, e.g., Glu542-Lys576’ salt bridges detected with EDC are compatible with conformation #11, and DSBU linkage between residues Tyr545 and Tyr638 is only possible in homodimer #9 (Fig. 5A and Supplementary Table 6).

Our live-cell imaging studies have revealed that dimeric GR might be an intermediate state towards transcriptionally active tetramers bound to target DNA sequences (Presman *et al*, 2016; Paakinaho *et al*, 2019; Johnson *et al*, 2021), also in line with the current SPR results. Although more speculative, it is possible to generate tentative models of DNA-bound GR tetramers that satisfy the constraints derived from current structure-function information. First, we reasoned that since tetramers but not higher order multimers are detected on chromatin, none of the known intermonomer “anchors”, Tyr545 and Ile628, could be available for protein-protein interactions in active tetramers. In other words, all GR-LBD “valences” should be satisfied in *bona fide* GR tetramers. Since the Ile628 side chain is exposed in both conformations with opposing Tyr545 rings (#6 and #11), different combinations with homodimers #9 and #10 were tested first. Indeed, different trimeric and tetrameric arrangements could be envisioned, eventually upon replacing the “true” crystal monomer by a close docking solution to avoid intermonomer clashes (Supplementary Figs. 5E-J). These hypothetical multimeric arrangements reconcile the role of residues Tyr545 and Ile628 for receptor multimerization. The additional interactions predicted in these multimeric arrangements, in addition to DBD-DBD interactions (Presman *et al*, 2016) and condensation provided by the NTD (Frank *et al*, 2018; Stortz *et al*, 2020) would overcome the energy loss due to Tyr545→Ala or Ile628→Ala exchanges, explaining why variants Y545A and I628A are still tetrameric on chromatin (Fig. 6D). However, the triple mutant (A458T, Y545A, I628A) did not bind the array, indicating that simultaneous elimination of the Tyr545 and Ile628 valences would generate a well-folded, but fully inactive variant.

In summary, we have explored the multivalency of GR-LBD and have associated experimentally observed homodimers to specific pathophysiologically relevant states of this NR. Homodimers with significantly larger interface surfaces are uniquely linked to antagonist-bound conformations and are likely to be found only in inactive / self-repressed states of the GR. Further, we provide evidence indicating that a mutant associated with GC resistance, p.Asp641Val, might form homo-hexamers on chromatin, and we present a 3D model of these hexamers. Finally, we have identified a previously unappreciated role of the Tyr545 phenolic ring for receptor homodimerization. Current structure-function information suggests that four different GR-LBD homodimers have roughly equal probabilities to be formed in vivo, which would in turn generate different tetrameric arrangements on chromatin. We are tempted to speculate that these individual conformations of tetrameric GR are associated with specific transcription programs. An inspection of predicted GR-LBD tetramers suggests mechanisms to select these different conformations. For instance, a peptide corresponding to the third LXXLL motif of SRC2/TIF2 would severely clash with a neighboring monomer in some arrangements. Thus, it is possible that different coregulators, or even different LXXLL motifs within a given coactivator select or induce specific NR quaternary arrangements. TNFα-induced modulation of the GR interactome, in particular weakened interactions with p300, and its implications for GR transcriptional output (Dendoncker *et al*, 2019) are in line with this suggestion. Future investigations should verify the validity of this hypothesis and establish whether specific GR conformations are associated with unique expression patterns. The ability to promote or stabilize these specific multimeric states might be an essential step towards the development of novel GCs with reduced side effects.

## Acknowledgments

We thank Prof. Erick Ortlund (Emory University) for providing the plasmid for ancGR2-LBD expression, Arnold T. Hagler for his comments on the manuscript, and Ildefonso Cases (CABD) for his help and input with some code. We thank ALBA-Cells synchrotron Xaloc team for beamline support.

## Author contributions

P.F.-P. and E.E.-P. conceived and designed the initial project and supervised its overall execution, provided financial support, and share overall responsibility for the presented results and final approval of the article. A.J.P., A.A.M., M.A.M. and R.A. expressed and purified recombinant proteins. M.A.M., A.A.M., and A.J.P. performed crystallization trials. A.A.M, J.F.D. and R.A. performed site-directed mutagenesis and protein analyses. I.N.B., A.R.M., and E.E.-P. performed and interpreted sequence and SCA analyses. J.F.R. and A.J.P. performed and interpreted docking, NIP and ODA calculations. T.T., R.L.S., and T.A.J. generated GR-FL mutants. G.F. and D.M.P. performed and analyzed N&B experiments. A.J.P., P.F.-P., and E.E.-P. performed SPR experiments and interpreted data. A.J.P., R.A., P.F.-P., and E.E.-P. performed MS experiments and interpreted results. A.J.P., P.F.-P., and E.E.-P. performed X-ray crystallography experiments, interpreted diffraction data and solved and refined 3D structures. T.T., R.L.S., T.A.J., A.V.F., C.C., and P.P. contributed tools. D.M.P., and G.L.H. supervised cell experiments. A.J.P., P.F.-P., and E.E.-P. drafted the article. A.R.M, J.F.R., D.M.P., G.L.H., P.F.-P., and E.E.-P. critically reviewed the manuscript. All authors discussed the results and commented on the manuscript.

## Data and materials availability

The atomic coordinates and structure factors have been deposited in the Protein Data Bank (PDB; www.rcsb.org) and the accession codes assigned are xxxx (C2), wwww (P3_1_), yyyy (P6_1_), zzzz (I4_1_22), and vvvv (I4_1_32). The PDB accessibility has been designed “for immediate release on publication”. The mass spectrometry proteomics data have been deposited to the ProteomeXchange Consortium via the PRIDE partner repository with the dataset identifier PXD028039. Bioinformatics data and code for the clustering, and sector analyses are available at https://github.com/ibn90/GRPROJECT2021. The authors declare that all relevant data are available upon request.

## Abbreviations

AF: Activation function

AR: Androgen receptor

ASA: Accessible surface area

ASU: Asymmetric unit

BF-3: Binding function-3

BSA: Buried surface area

CHAPS: 3-[(3-cholamidopropyl) dimethylammonio]-1-propanesulfonate

DBD: DNA binding domain

DEX: Dexamethasone

EDC: 1-ethyl-3-(3-dimethylaminopropyl)carbodiimide

EM: Electron microscopy

ER: Estrogen receptor

FL: Full-length

GC: Glucocorticoid

GR: Glucocorticoid receptor

GRE: Glucocorticoid response elements

LBD: Ligand-binding domain

LBP: Ligand-binding pocket

MS: Mass spectrometry

MR: Mineralocorticoid receptor

N&B: Number and brightness

NIP: Normalized interface propensity

NTD: N-terminal domain

ODA: Optimal docking area

PDB: Protein Data Bank

PR: Progesterone receptor

RMSD: Root-mean-square deviation

SHP: Small heterodimer partner

SPR: Surface plasmon resonance

WT: Wild type

## Methods

### Peptides and proteins

A peptide corresponding to residues Gln18-Lys27 of SHP/NR0B2 (box 1 motif; NH2-Q_18_GAASRPAILYALLSSSLK_27_-OH) was custom-synthesized at Pepmic. Recombinant ancGR2-LBD (corresponding to residues 529 to 777 of the human receptor) cloned into a pMALCH10T vector was expressed as fusion protein with an N-terminal maltose-binding protein (MBP) and a hexahistidine tag and purified to homogeneity using standard chromatographic procedures (Weikum *et al*, 2017b).

### Crystallization and structure determination

Purified, concentrated DEX-bound ancGR2-LBD was combined with a 3-fold molar excess of SHP peptide and incubated for one hour at RT. Drops of the ancGR2-LBD-SHP mixture were equilibrated against 0.1 M Tris-HCl, pH 8.0, 0.2 M sodium chloride, 2.0 M ammonium sulfate (P3_1_, I4_1_22 and I4_1_32 crystals); 0.1 M PIPES, pH 7.0, 0.1 M ammonium acetate, 2.5 M sodium formate (P6_1_ crystals); or 85 mM sodium cacodylate trihydrate, pH 6.5, 0.17 M sodium acetate trihydrate, 25.5% (w/v) PEG8000, 15% (v/v) glycerol (C2 crystals) using the sitting drop vapor-diffusion method. Diffraction data were collected at 100 K at the ALBA-CELLS synchrotron and processed using MOSFLM (http://www.mrc-lmb.cam.ac.uk/harry/mosflm/) and CCP4 (http://www.ccp4.ac.uk/). The crystal structures were solved and refined using MOLREP, REFMAC5 and COOT from the CCP4 package. Crystal packing was analyzed using PISA (http://www.ebi.ac.uk/), model quality was assessed with MolProbity (http://molprobity.biochem.duke.edu/), and figures were prepared with PyMOL (http://www.pymol.org).

### Surface plasmon resonance (SPR) analyses

SPR analyses were performed at 25°C in a BIAcore T200 instrument (GE Healthcare). Highly purified, DEX-bound recombinant WT ancGR2-LBD and its Y545C and Y545A mutants were diluted in 10 mM sodium acetate, pH 5.0 and directly immobilized on CM5 chips (GE Healthcare) by amine coupling at densities between 300 and 400 resonance units (RU). As a reference, one of the channels was also amine-activated and blocked in the absence of protein. The running buffer was 50 mM HEPES, pH 7.2, 50 mM Li_2_SO4, 5% glycerol, 1 mM dithiothreitol (DTT), 50 µM DEX. Sensorgrams were analyzed with the BIAcore T200 Evaluation software 3.0 and fitted according to the Langmuir 1:1 and multisite models.

### Crosslinking experiments

Purified recombinant ancGR2-LBD (33 µM) was incubated with 100-fold molar excess of crosslinkers 1-ethyl-3-(3-dimethylaminopropyl)carbodiimide hydrochloride (EDC, Pierce) or disuccinimidyl dibutyric acid (DSBU) for 1-2 hours at 37 °C following the manufacturer’s instructions. In some experiments the Y545C variant was incubated after affinity purification for 30 min at room temperature without further treatment. Samples of the reaction mixtures were boiled in the presence of Laemmli sample buffer, either reducing (EDC- and DSBU-crosslinked proteins) or non-reducing (in the case of the Y545C variant) and resolved by SDS-PAGE.

### Nano-LC-MS/MS mass spectrometry

CBB-stained bands corresponding to monomeric, dimeric and tetrameric GR-LBD after crosslinking with EDC or DSBU were excised from the gels and subjected to in-gel digestion following standard protocols. Briefly, excised bands were reduced (10 mM DTT) in 50 mM bicarbonate buffer, pH 8.0, for 45 min at 56 °C, alkylated (50 mM iodoacetamide in 50 mM ammonium bicarbonate buffer for 30 min at 25 °C) and digested with either trypsin alone or followed by GluC treatment, or with chymotrypsin overnight at 37 °C in 100 mM ammonium acetate buffer, pH 8.0. (Sequencing-grade endoproteases were from Promega). In the case of the Y545C mutant, proteins in the excised bands were directly alkylated without previous DTT treatment to prevent reduction of the Cys545-Cys545’ disulfide bridge.

Tryptic peptides were diluted in 1% formic acid and loaded onto a 180 µm x 20 mm C18 Symmetry trap column (Waters) at a flow rate of 15 µl/min using a nanoAcquity Ultra Performance LCTM chromatographic system (Waters). Peptides were separated using a C18 analytical column (BEH130 C18, 75 mm x 25 cm, 1.7 μm; Waters) with a 120-min run, comprising three consecutive linear gradients: from 1 to 35% B in 100 min, from 35 to 50% B in 10 min and from 50 to 85% B in 10 min (A= 0.1% formic acid in water, B= 0.1% formic acid in CH_3_CN). The column outlet was directly connected to an Advion TriVersa NanoMate (Advion) fitted on an LTQ-FT Ultra mass spectrometer (Thermo), which was operated in positive mode using the data-dependent acquisition mode. Survey MS scans were acquired in the FT-ICR cell with the resolution (defined at 400 m/z) set to 100,000. Up to six of the most intense ions per scan were fragmented and detected in the linear ion trap. The ion count target value was 1,000,000 for the survey scan and 50,000 for the MS/MS scan. Target ions already selected for MS/MS were dynamically excluded for 30 s. Spray voltage in the NanoMate source was set to 1.70 kV. Capillary voltage and tube lens on the LTQ-FT were tuned to 40 and 120 V, respectively. The minimum signal required to trigger MS to MS/MS switch was set to 1,000 and activation Q value was set at 0.25. Singly charged precursors were rejected for fragmentation.

### Differential scanning fluorometry

Thermofluor experiments were performed in an iQ5 Multicolor Real Time PCR Detection System (BIO-RAD) using 96-well plates (Hard-Shell® High-Profile Semi-Skirted PCR Plate, BIO-RAD) and a 25-µL total volume for each reaction. Melting curves were acquired from eight replicates to determine the average melting temperature (Tm). GR-LBD samples (0.5 mg/mL) were prepared in 20 mM HEPES, pH 8.0, 200 mM NaCl, 10% glycerol, 50 mM imidazole, 1 mM DTT, 50 µM DEX, and centrifuged 5 min at 14,000 rpm immediately before measurements. SYPRO Orange dye (Sigma-Aldrich) was firstly prepared at 80× in the same buffer, starting from a 5,000× commercial dilution. The final concentration of the dye in each well was 5×. The plates were sealed with optical-quality sealing film (Microseal® B Seals, BIO-RAD) and centrifuged at 2,000×g for 30 s. Samples were equilibrated for 60 s and analyzed using a linear gradient from 16 to 95 °C with increments of 1 °C/min, recording the SYPRO orange fluorescence throughout the gradient using the iQ5 Optical System Software 2.0. Values were fitted using the online tool JTSA with the four-parameter logistic equation, and the calculated fluorescence midpoints were compared with an unpaired t-test for equal variances using GraphPad Prism 8.

### Subcellular localization and number and brightness (N&B) analysis

pROSA-GFPmGR expresses GFP-tagged mouse GR under the CMV promoter. The plasmid also contains homologous recombination arms for potential integration into the GT(Rosa)26Sor locus (Paakinaho *et al*, 2019). All mutations were generated with the QuikChange II XL Site-Directed Mutagenesis Kit (Stratagene) according to the manufacturer’s instructions.

Mammary adenocarcinoma 3617-derived GR^null^ cells (Paakinaho *et al*, 2019) were grown in Dulbecco’s modified Eagle’s medium (DMEM, Invitrogen) supplemented with 5 μg/ml tetracycline (Sigma-Aldrich), 10% fetal bovine serum (Gemini), sodium pyruvate, non-essential amino acids, and 2 mM L-glutamine maintained in a humidifier at 37 °C and 5% CO_2_. This cell line contains a tandem array (∼200 copies) of a mouse mammary tumor virus long terminal repeat, Harvey viral ras (MMTV-v-Ha-ras) reporter integrated into chromosome 4, which can be directly visualized in living cells as a localized domain if bound to a fluorescently labelled protein63. Prior to DEX treatment, cells were seeded to 2-well Lab-Tek chamber slides (Thermo Fisher, Waltham, MA, USA) and incubated for at least 18 h in DMEM medium containing 10% charcoal-stripped FBS (Life Technologies) and 2 mM L-glutamine. Cells were transiently transfected with the indicated pROSA-GFPmGR mutants using jetOPTIMUS™ reagent (PolyPlus) according to the manufacturer’s instructions.

Images were taken at the CCR, LRBGE Optical Microscopy Core facility in a LSM 780 laser scanning microscope (Carl Zeiss, Inc.) equipped with an environmental chamber. Cells were imaged from 20 min up to a maximum of 2 hours after DEX addition. We used a 63× oil immersion objective (NA = 1.4). The excitation source was a multi-line Ar laser tuned at 488 nm. Fluorescence was detected with a GaAsP detector in photon-counting mode.

N&B measurements were done as previously described (Presman *et al*, 2016). For each studied cell, a single-plane stack of 150 images (256 x 256 pixels) were taken in the conditions mentioned above, setting the pixel size to 80-nm and the pixel dwell time to 6.3 µs. In all cases, we discarded the first 10 images of the sequence to reduce overall bleaching. The frame time under these conditions is 0.97 s, which guarantees independent sampling of molecules according to previously reported FCS measurements (Mikuni *et al*, 2007). Each stack was further analyzed using the N&B routine of the SIMFCS 2.0 software developed at the Laboratory for Fluorescence Dynamics (UCI). In this routine, the average fluorescence intensity () and its variance (σ2) at each pixel of an image are determined from the intensity values obtained at the given pixel along the image stack. The apparent brightness (B) is then calculated as the ratio of σ2 to while the apparent number of moving particles (N) corresponds to the ratio of to B (Digman *et al*, 2008). Classification of pixels according to their intensity values allows to easily split nucleus and array for further analysis. Selection of cells for analysis followed these criteria. (1) In the case of stimulated cells, an accumulation of signal at the array must be visible. (2) The average apparent number of molecules (N) in the nuclear compartment must have a range of 3-18 in all cases, (3) no saturation of the detector at any pixel (N < 60), and (4) bleaching cannot exceed 5-10%. In a previous work it has been demonstrated that B is equal to the real brightness ε of the particles plus one (Digman *et al*, 2008). Therefore, ε at every pixel of images can be easily extracted from B measurements. Importantly, this analysis only provides information regarding the moving or fluctuating fluorescent molecules since fixed molecules (relative to our frame time) will give B values equal to 1. The experiments were independently repeated two times for each treatment/condition.

### Bioinformatics analysis of the impact of GR point mutations

We estimated the impact of the generated mutations on the overall protein stability with the FoldX empirical force (http://foldxsuite.crg.eu/), which has an estimated error of ∼0.7 kcal/mol. Ten iterations were conducted for each mutation, and later averaged. Free energy differences between mutant and WT proteins (ΔΔG) < 1 kcal/mol were considered not significant, those between 1 and 2, 2 and 4, and > 4 kcal/mol as slightly, mildly, and strongly destabilizing, respectively.

### Sequence analyses

The sequence of human GR-LBD was used as query against the whole NR database, from which we selected ∼880 sequences and included the ancestral GR(Weikum *et al*, 2017b). We also downloaded the sequences corresponding to the LBD regions from a representative fraction of proteomes at PFAM rp55 (PF00104_rp55). (Note that the sequences included in the PFAM alignment are truncated, as they lack for instance the non-conserved F-domain). We followed three different approaches to ensure sequence and alignment diversity and thus stability of the analyses. First, we aligned the ∼880 sequences to a structure-based profile from entries 5UFS (GR-LBD) and 5JJM (AR-LBD). The resulting alignment, 880_aln, was used to run pySCA in addition to the original SCA5 method. Secondly, we aligned our ∼880 sequences to a profile generated from the PF00104_rp55 removing fragments to generate the 840_aln. Finally, we used the PFAM alignment as retrieved from the PFAM database, which contains ∼13,000 sequences (PF00104_rp55).

### Multiple correspondence analysis (MCA): identifying specificity-determining positions

Positions differentially conserved within protein subfamilies (termed “specificity determining positions”, SDPs) are related to functional specificity (e.g., binding of different cofactors; for a review see Pazos & Bang, 2008). Recent methods based on multiple correspondence analysis (MCA) have allowed identification of their subtle patterns of conservation within large protein families (Rojas *et al*, 2012). We have performed both supervised and unsupervised runs of the S3DET method (http://csbg.cnb.csic.es/JDet/) on 840_aln using default parameters to maintain a large sequence diversity.

### Statistical coupling analysis (SCA)

We have used the SCA5 8/2011 version(Halabi *et al*, 2009) with the three different alignment versions given above, and the updated version pySCA67 with 880_aln. Each alignment produced unique sets of residues termed Sca5_880, Sca5_840, and Sca5_onPFAMrp55, respectively. Next, we labeled residues in the three different sectors that emerged as outputs of the program with letters A, B, and C, following their order of appearance. The stability of the identified sectors was assessed with a statistical test based on hypergeometric calculations of the groups of residues belonging to given sectors between pairs of alignments. P-values were adjusted using false discovery rate (FDR). Next, specific residues from the significant sectors were extracted and selected according to their rank. For instance, if a particular residue appears only in one sector on a low-ranking pair of alignments (e.g., rank 19, with a borderline p-value) this residue will not be selected as part of a sector. On the contrary, if a residue appears in high-ranking pairs, it will be retained. Residues termed as “A” and “B” appeared to be equivalent in different pairs of alignments, so they were assigned to the class sector 2, while residues belonging to the “C” group were stable, and therefore assigned to class sector 1.

### Clustering of interaction surfaces in GR-LBD dimers

We grouped the interaction surfaces observed in the 20 GR-LBD homodimers by a hierarchical clustering analysis, using *ad-hoc* R scripts. For each dimer, the interaction surface was defined as the set of solvent-exposed residues in the monomers (i.e., residues with > 25% relative accessible surface area, ASA) that became buried (< 25% relative ASA) in the corresponding dimer. Relative ASA were calculated using ICM (Molsoft LLC). Then, we computed the center of coordinates of the residues forming each interaction interface. To compare two pairs of homodimers, we computed the Euclidean distances between the centers of their interaction surfaces after superimposition on a common monomer. The 40×40 distance matrix representing the distances between all pairs of interaction surfaces was used to perform hierarchical clustering with Ward’s method. Finally, the dendrogram generated from this analysis was sorted in order of increasing distance.

### Docking experiments and analysis

Homodimeric models of GR-LBD were built using pyDock docking and scoring method68. First, protein models were prepared by removing all cofactors and heteroatoms, and missing side chains were modeled with SCWRL 3.0 (Canutescu et al, 2003). Then, the Fast Fourier Transform (FFT)-based docking programs FTDock70 (with electrostatics and 0.7-Å grid resolution) and ZDOCK 2.1 (Chen *et al*, 2003) were used to generate 10,000 and 2,000 rigid-body docking poses, respectively. These were merged in a single pool for subsequent pyDock scoring68, based on energy terms previously optimized for rigid-body docking. The pyDock binding energy is basically composed of accessible surface area (ASA)-based desolvation, Coulombic electrostatics and VdW energy terms. Electrostatics and VdW contributions were limited to −1.0/+1.0 and 1.0 kcal/mol for each inter-atomic energy value, respectively, to avoid excessive penalization from possible clashes derived from the rigid-body approach.

### Predicted dimer interfaces

Optimal docking areas (ODA) per surface-exposed protein residues were obtained by computing surface patches with optimal desolvation energy based on the selection of low-energy docking regions generated from each surface residue (Fernandez-Recio *et al*, 2005). ODA hot spots (residues with low ODA values, usually less than −10.0 kcal/mol) indicate regions with favorable desolvation energy upon interaction with a partner protein.

From the resulting docking poses, normalized interface propensity (NIP) values were obtained for each residue with the built-in patch module of pyDock, implementing the pyDockNIP algorithm (Grosdidier & Fernández-Recio, 2008). A NIP value of 1 indicates that the corresponding residue is involved in all predicted interfaces of the 100 lowest energy docking solutions, while a value of 0 means that it appears as expected by random chance. Finally, a negative NIP value implies that the residue appears at the low-energy docking interfaces less often than expected by random chance. Usually, residues with NIP ≥ 0.2 are considered as hot-spot residues when using FTDock.

### Energetic characterization of GR dimers

The binding energy of the different crystal dimers was computed with the pyDock bindEy module, using the same scoring function as in docking (Cheng *et al*, 2007).

## Supplementary figure legends and Tables

**Fig. S1.**
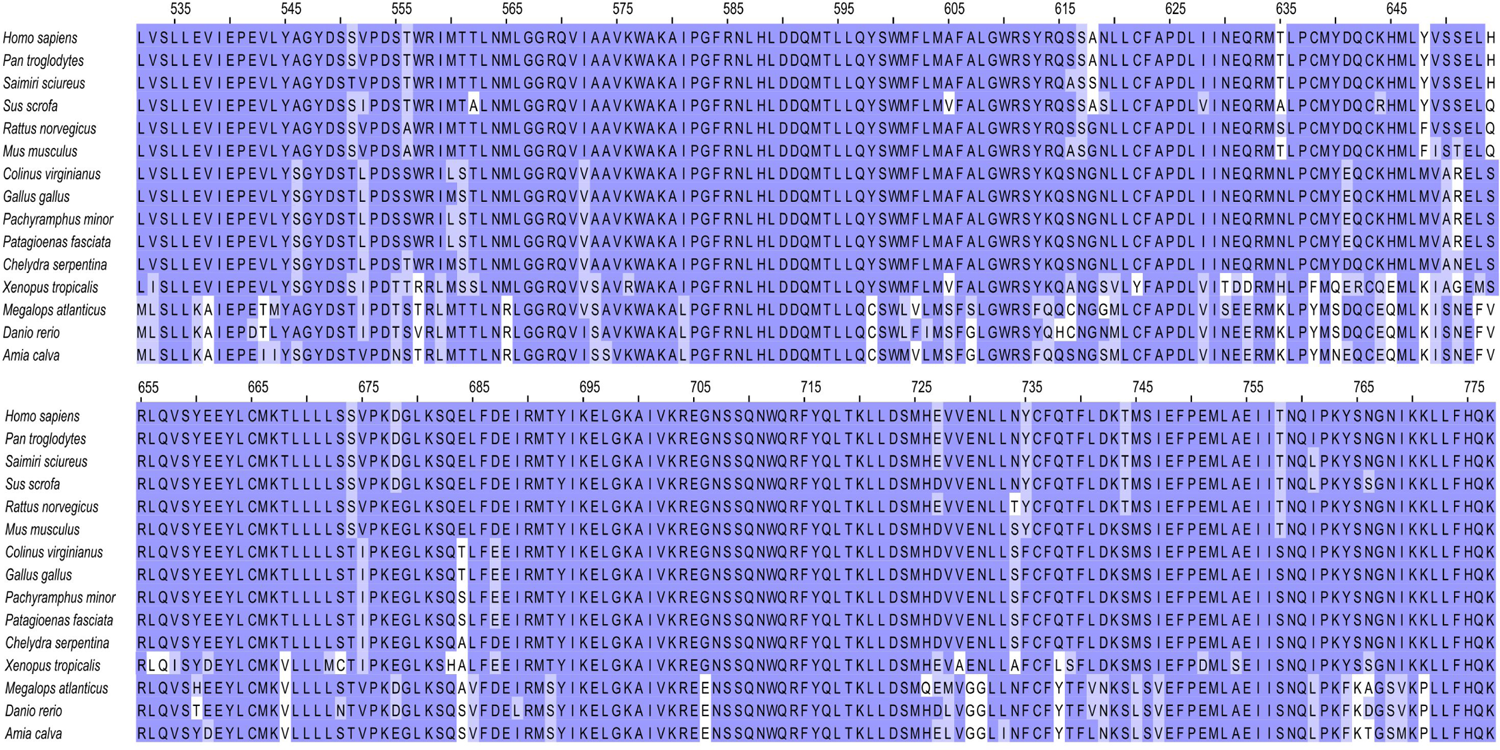
legend. Partial sequence alignment of GR-LBD from selected species. GR homologs were identified and aligned using PSI-BLAST (https://blast.ncbi.nlm.nih.gov/Blast.cgi?PAGE_TYPE=BlastSearch&PROGRAM=blastp&BLAST_PROGRAMS=psiBlast). To provide a representative view of LBD conservation across vertebrates, sequences of mammalian (human, chimpanzee, squirrel monkey, pig, rat, and mouse), avian (chicken, quail, becard, and pigeon), reptile (turtle), amphibian (frog) and fish GR (tarpon, zebrafish, and dogfish) were included in the alignment. Residues conserved in hGR and other homologs are shaded purple, and conservative replacements are shaded light violet.

**Fig. S2.**
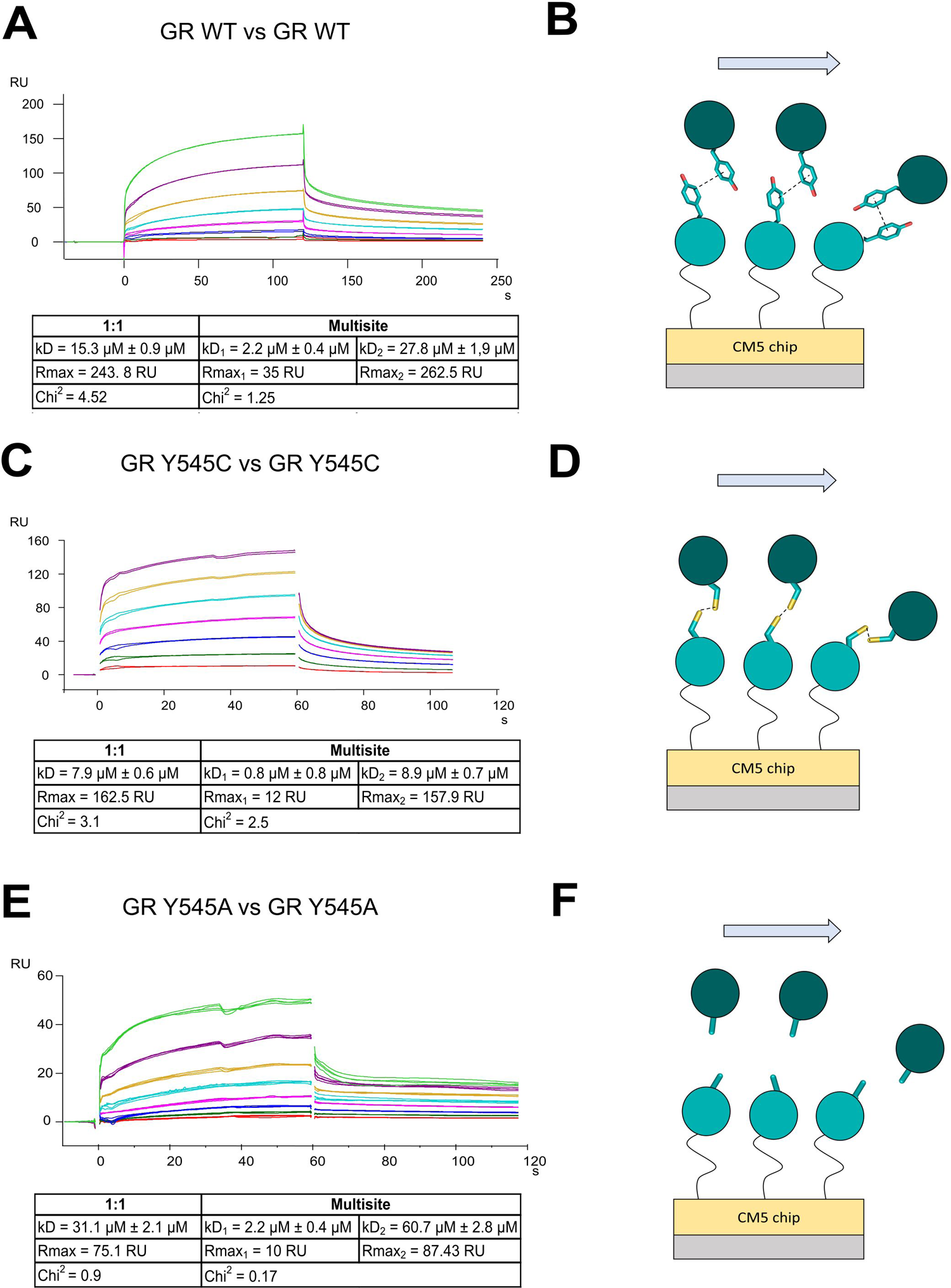
legend. SPR analysis of protein-protein interactions between wild-type GR-LBD and its Tyr545 mutants. SPR experiments were performed by running increasing concentrations (0.2, 0.4, 0.8, 1.6, 3.12, 6.25, 12.5 and 25 µM) of DEX-bound WT GR-LBD (**A**), GR-LBD Y545C mutant (**C**), or GR-LBD Y545A mutant (**E**) over the same immobilized protein. SPR sensorgrams corresponding to experiments conducted in duplicate are shown in all cases. A schematic representation of the interactions between soluble analyte and chip-immobilized molecules are depicted in panels **B**, **D** and **F**, respectively. Tables below panels **A, C** and **E** summarize major parameters (maximum response (R_max_), dissociation constant (k_D_), and fitting error (Chi^2^)) derived from the fitting of self-association sensorgrams according to either 1:1 or multisite models.

**Fig. S3.**
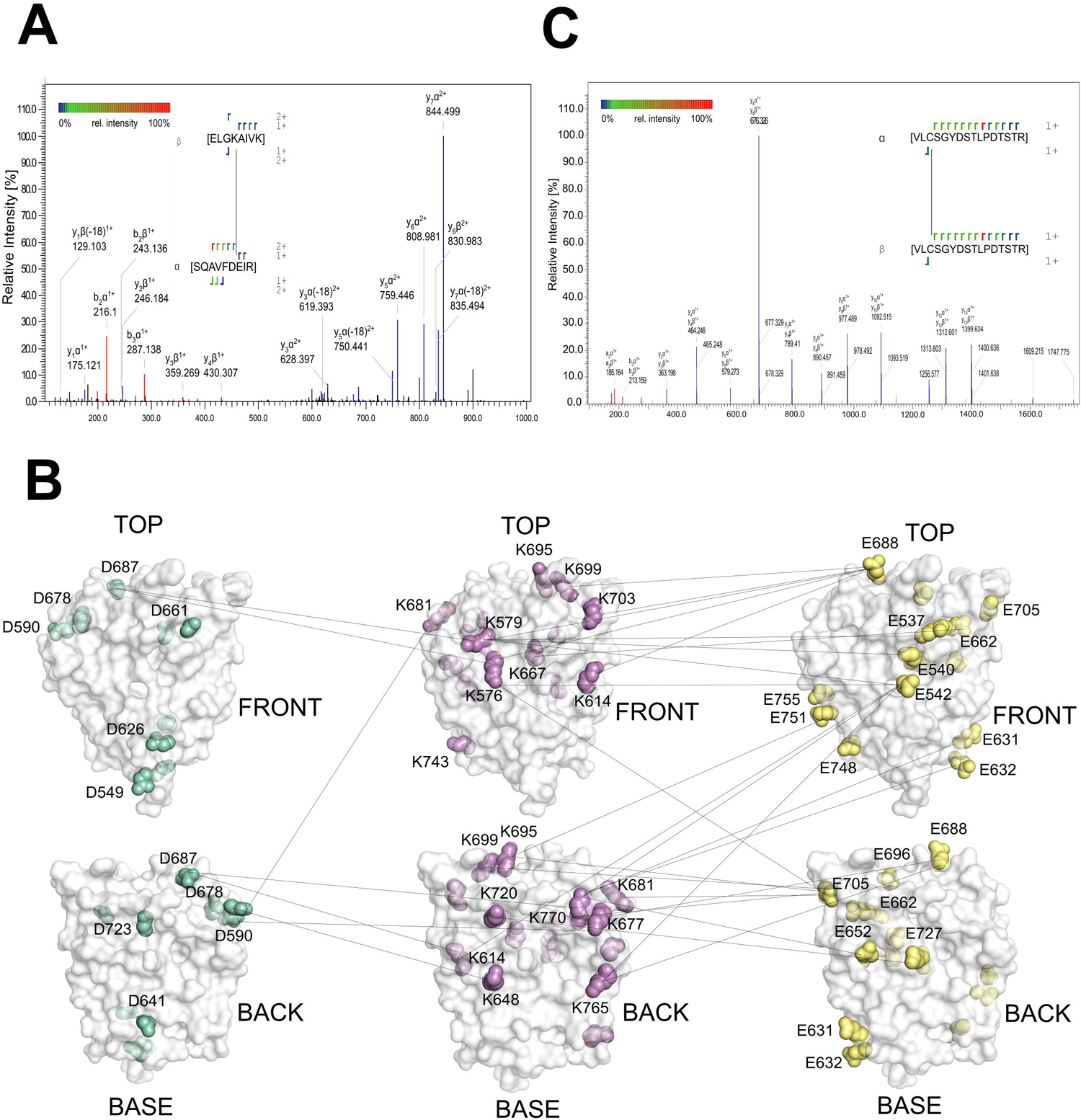
legend. Mass spectrometric verification of cross-links between GR-LBD molecules in solution. **A**, Representative MS/MS spectra identifying EDC-crosslinked peptides between Glu688 and Lys699, corresponding to the top-to-top conformation (#20). **B,** Summary of all EDC-crosslinks identified between Lys-Asp and Lys-Glu residue pairs, mapped on the 3D structure of GR-LBD. The central panels show the surface-exposed lysines, while aspartate and glutamate residues are highlighted in the left and right panels, respectively. Note that the uneven distribution of charged residues across the domain surface, in particular of lysines, only allows demonstration of a subset of possible homodimeric conformations populated in solution. **C**, Representative MS/MS spectra identifying crosslinked peptides formed by disulfide bridges between Cys545-Cys545’ residues in the Y545C variant of GR-LBD.

**Fig. S4.**
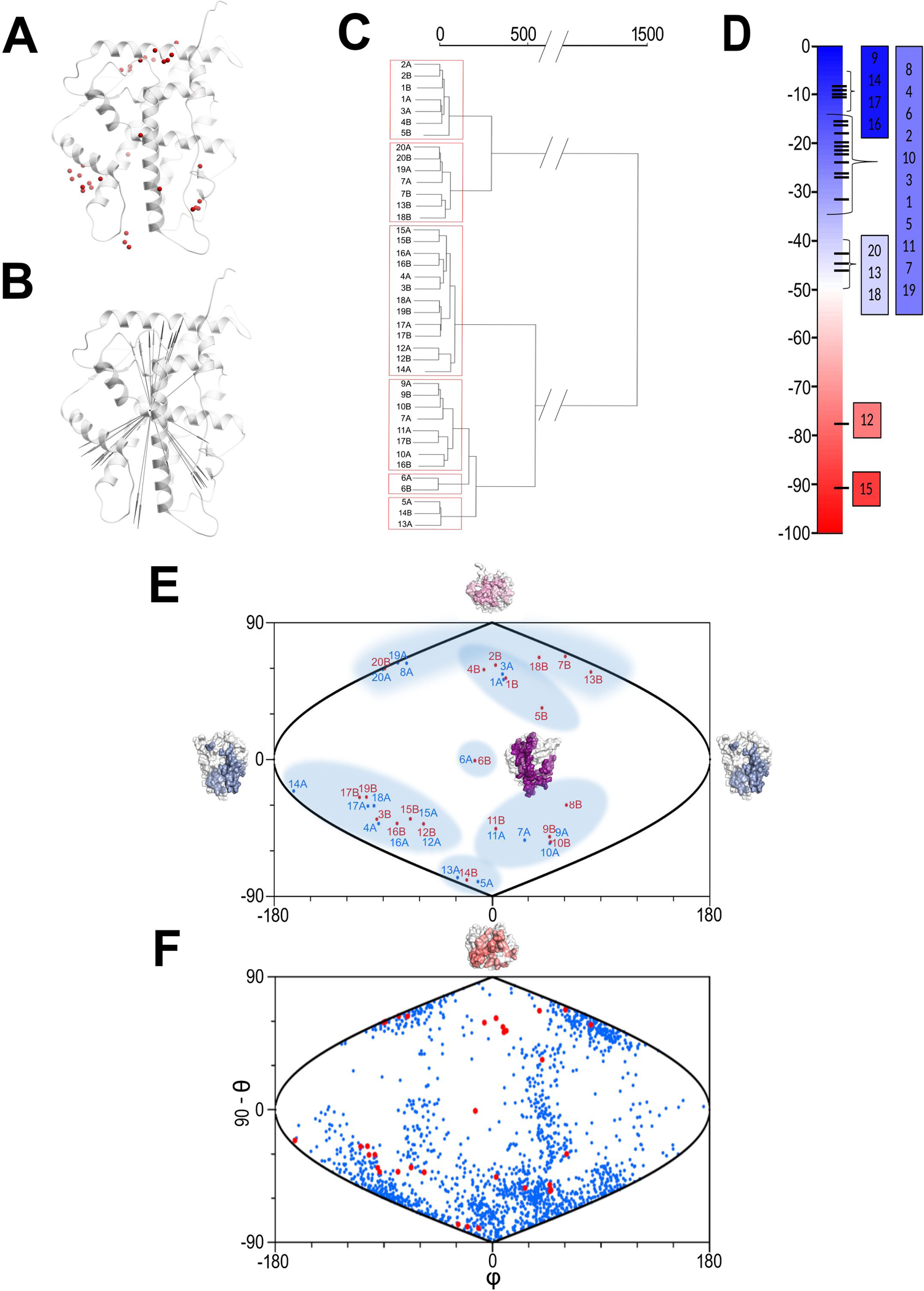
legend. Distribution and clustering of homodimerization interfaces in GR-LBD. The centers of coordinates of the interaction surfaces in the 20 topologically distinct GR-LBD homodimers are schematically represented as dots (**A**) or as vectors drawn from the center of coordinates of a common monomer to the given interface (**B**). **C**, Dendrogram generated from the hierarchical clustering of all GR-LBD homodimers, based on the Euclidean distances between the centers of coordinates of the interaction surfaces. Each interaction surface is named according to the dimer number (#1-#20) and the chain ID of the monomer. Note that the clustering analysis clearly separates the interaction surfaces in six clusters. **D**, Binding energy of the GR-LBD homodimers, as calculated with pyDock. **E**, A sinusoidal equal-area projection of the interaction surfaces in the 20 GR-LBD dimers, represented as spherical coordinates from the vectors defined in panel B. Clusters obtained as described above are shaded blue. **F**, Representation in spherical coordinates of the interaction surfaces in the top 1,000 docking solutions of monomeric GR-LBD on itself (blue dots, results obtained using coordinates form PDB entry 5UFS). For comparison, the interaction surfaces for the 20 topologically distinct crystal homodimers are shown as red dots.

**Fig. S5.**
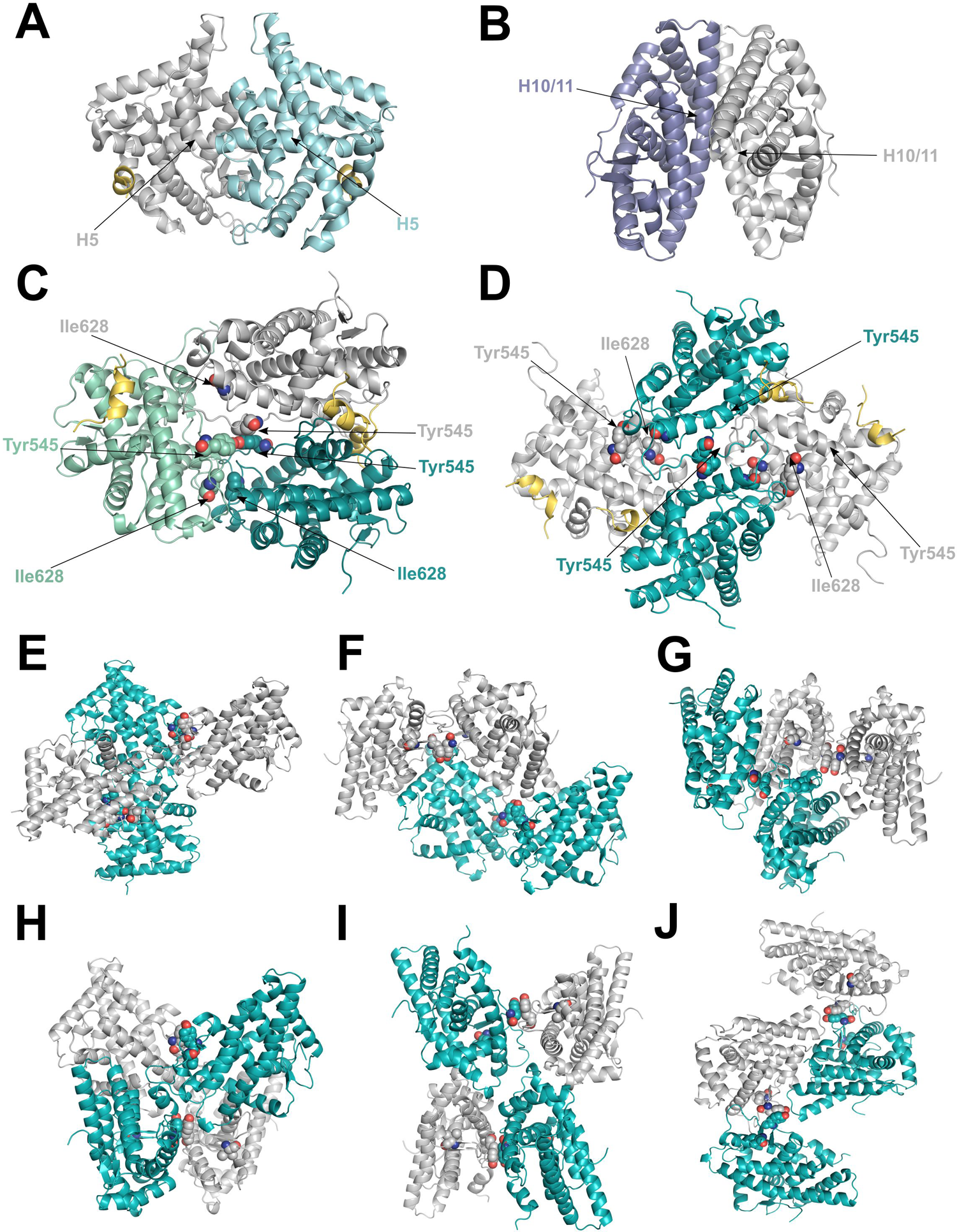
legend. Possible conformations of multimeric GR-LBD. **A**, Three-dimensional structure of dimeric AR-LBD (PDB 5JJM), a non-canonical prototype of NR dimerization. The two monomers are shown as cartoons, colored gray and light cyan, respectively. The interacting helices H5 from both monomers are labeled. **B**, Three-dimensional structure of ERα-LBD dimer (PDB 1ERE), representing the canonical conformation observed in NR homo- and heterodimers. The two monomers are shown as cartoons, colored slate blue and gray, respectively. The interacting helices H10/11 from both monomers are labeled. **C**, GR-LBD trimer generated by docking dimers #6 and #10. **D**, GR-LBD tetramer generated by combining dimers #11 and #14. **E-J**, Putative alternative conformations of tetrameric GR-LBD generated by docking dimer #11 onto itself. The major interface residues Tyr545 and Ile628 are given as spheres in panels **C** - **J**. Note that there are no “free valences” (exposed Tyr545 / Ile628 side chains) in these multimeric arrangements.

**Table S1.** legend. Summary of X-ray diffraction data, refinement statistics and model quality.

|  | GR WT 1 | GR WT 2 | GR WT 3 | GR WT 4 | GR WT 5 |
| --- | --- | --- | --- | --- | --- |
| **PDB code** | XXX | YYY | ZZZ | VVV | WWW |
| **Wavelength (Å)** | 0.9 | 0.9 | 0.9 | 0.9 | 0.9 |
| **Resolution range** | 43.42-2.25<br>(2.32-2.25) | 76.81-2.30<br>(2.42-2.30) | 94.81-2.46<br>(2.56-2.46) | 90.10-2.99<br>(3.18-2.99) | 72.80-3.36<br>(3.63-3.36) |
| **Space group** | C121 | P31 | P61 | I4<sub>1</sub>22 | I4<sub>1</sub>32 |
| **Cell dimensions**<br>**a, b, c (Å)**<br>**α, β, γ (°)** | 86.61, 52.44,<br>69.60<br>90.0, 116.62,<br>90.0 | 107.60,<br>107.60,<br>135.64<br>90.0, 90.0,<br>120.0 | 109.36,109.36,<br>137.77<br>90.0, 90.0,<br>120.0 | 180.20,<br>180.20,<br>169.18,<br>90.0, 90.0,<br>90.0 | 178.05,<br>178.05,<br>178.05<br>90.00, 90.00,<br>90.00 |
| **Total reflections** | 34,022<br>(1,954) | 156,013<br>(22,547) | 94,861<br>(11,087) | 223,344<br>(40,285) | 82,328<br>(16,909) |
| **Unique reflections** | 12,530<br>(907) | 73,686<br>(10,882) | 33,009<br>(3,799) | 23,704<br>(4,229) | 7,178<br>(1,431) |
| **Multiplicity** | 2.7 (2.2) | 2.1 (2.1) | 2.9 (2.9) | 9.4 (9.5) | 11.5 (11.8) |
| **Completeness (%)** | 93.8 (75.5) | 94.4 (95.6) | 96.8 (98.5) | 100.0 (100.0) | 100.0<br>(100.0) |
| **Mean I/sigma(I)** | 11.1 (1.8) | 5.9 (1.4) | 7.4 (0.7) | 6.6 (2.2) | 10.8 (1.7) |
| **Wilson B-factor** | 44.7 | 45.0 | 57.5 | 73.5 | 141.0 |
| **R-meas** | 0.069 (0.697) | 0.125 (0.987) | 0.074 (1.050) | 0.307 (1.701) | 0.108 (1.762) |
| **R-pim** | 0.039 (0.439) | 0.079 (0.641) | 0.051 (0.725) | 0.149 (0.815) | 0.031 (0.505) |
| **CC1/2** | 0.997 (0.654) | 0.986 (0.314) | 0.994 (0.317) | 0.982 (0.386) | 1.000 (0.561) |
| **Reflections used in refinement** | 12,018 | 69,901 | 31,272 | 22,492 | 6,809 |
| **Reflections used for R-free** | 510 (4.1%) | 3,765 (5.1%) | 1,692 (5.1%) | 1,194 (5.0%) | 344 (4.8%) |
| **R-work** | 0.192 | 0.177 | 0.216 | 0.242 | 0.199 |
| **R-free** | 0.245 | 0.220 | 0.247 | 0.280 | 0.288 |
| **Total number of atoms** | 2,146 | 8280 | 4,190 | 6,109 | 2,111 |
| **RMS (bonds)** | 0.007 | 0.010 | 0.007 | 0.006 | 0.006 |
| **RMS (angles)** | 1.565 | 2.016 | 1.655 | 1.555 | 1.727 |
| **Ramachandran favored (%)** | 96.00 | 95.93 | 93.16 | 89.38% | 84.55 |
| **Ramachandran allowed (%)** | 100.0 | 99.3 | 99.0 | 98.9% | 95.9 |
| **Clashscore** | 12.86 | 13.16 | 9.36 | 27.61 | 29.6 |
| **Average B-factor** | 54.0 | 50.0 | 68.0 | 67.0 | 142.0 |

**Table S2.** legend. Summary of GR-LBD structures reported to date. PDB entries are ordered from lowest (monoclinic, C2) to highest (cubic, I4_1_32) space group symmetry. Major crystal parameters are given, along with the identity of LBP-bound compounds, peptides occupying the AF2 site, and other co-crystallized small-molecule compounds. ancGR, ancient GR; the sequences of ancGR1 and 2 are given in Fig. 1C. Other engineered variants of human GR (hGR; UniProt entry P04150) are named according to the first deposited structure. The mutations introduced in these variants are: hGR_1M2Z: F602S; hGR_3CLD: F602Y, C638G; hGR_1NHZ: F602S, C638D; hGR_3H52: F602S, C638D, E684A, E688A, W712S; hGR_4CSJ: V571M, F602S, C638D; hGR_5NFT: V571M, F602S, C638D, E684A, W712S; hGR_4P6X: F602A, C622Y, T668V, S674T, V675I, E684A, E688A; and hGR_4P6W: F602A, C622Y, T668V, S674T, V675I, K699A, K703A. PDB entry 5UC3 features a dominant negative hGR variant, (L733K, N734P). The reported structures of mouse GR-LBD contain either the F602S mutation (3MNE), the double mutant (F602S, A605V) in 3MNO, or the triplet (A605V, V702A, E705G) in 3MNP. HetgaGRb refers to the GRb variant from the naked mole-rat, *Heterocephalus glaber*, with residues topologically equivalent to Phe602 and Val728 replaced by Ser and Asn, respectively. Other abbreviations: 1CA / 1TA, triamcinolone acetonide; 29M, non-steroidal GR antagonist; 8W5, budesonide; 8W8, indazole ether-based GR modulator AZD5423; ACE, acetyl group; B9Q, B9T, and B9W, oral GR modulators; BOG, b-octyl glucopyranoside; CL, chloride ion; CHAPS, 3-[(3-cholamidopropyl)dimethylammonio]-1-propanesulfonate; CV7, desisobuytyryl ciclesonide; DAY, deacylcortivazol; DEX, dexamethasone; DMS, dimethyl sulfoxide; DOC, desoxycorticosterone; E7T, GR agonist; EDO, 1,2-ethylenediol; FMT, formic acid; GOL, glycerol; GSK866, non-steroidal GR agonist; GW6, fluticasone furoate; HCY, cortisol; HEPES, 4-(2-hydroxyethyl)-1-piperazine ethanesulfonic acid; HEX, hexane 1,6-diol; HYC, hydrocortisone; JZN, D-prolinamide 11; JZS, alanine amide derivative; JZR, hexyl b-D-glucopyranoside; LSJ, dibenzoxepane sulfonamide; MOF, mometasone furoate; MPD, (4*S*)-2-methyl-2,4-pentanediol; NN7, non-steroidal GR modulator, suitable for inhalation; R8C, indazole ether-based non-steroidal GR modulator; RU-468, mifepristone; SCN, thiocyanate ion; TLA, tartaric acid.

| PDB code | Space group | Res. (Å) | Cell constants | Protein |
| --- | --- | --- | --- | --- |
| 5UFS | C2 | 2.12 | a = 87.0, b = 52.8, c = 126.0 Å, $\beta = 101.8^\circ$ | ancGR2 |
| 3K23 | C2 | 3 | a = 184.9, b = 66.0, c = 71.5 Å, $\beta = 103.6^\circ$ | hGR_3CLD |
| 3RY9 | C2 | 1.95 | a = 139.4, b = 49.1, c = 100.4 Å, $\beta = 105.8^\circ$ | ancGR1[1] |
| 5UC3 | C2 | 2 | a = 136.7, b = 88.3, c = 64.7 Å, $\beta = 110.8^\circ$ | hGR_5UC3 |
| This work | C2 | 2.15 | a = 91.1, b = 90.8, c = 157.2 Å, $\beta = 116.6^\circ$ | ancGR2 |
| 1NHZ | P21212 | 2.3 | a = 74.9, b = 109.8, c = 39.3 Å | hGR_1NHZ |
| 4LSJ | P21212 | 2.35 | a = 39.0, b = 139.4, c = 48.1 Å | hGR_3CLD |
| 6NWK | C2221 | 1.65 | a = 71.7, b = 96.4, c = 107.9 Å | ancGR2 |
| 6NWL | C2221 | 1.59 | a = 71.7, b = 96.4, c = 108.2 Å | ancGR2 |
| 1P93[3] | P31 | 2.7 | a = b = 127.4, c = 91.8 Å, $\gamma = 120^\circ$ | hGR_1NHZ |
| This work | P31 | 2.7 | a = b = 107.75, c = 135.8 Å, $\gamma = 120^\circ$ | ancGR2 |
| 3GN8 | P61 | 2.5 | a = b = 104.2, c = 144.2 Å, $\gamma = 120^\circ$ | ancGR2 |
| 4E2J | P61 | 2.5 | a = b = 104.4, c = 143.9 Å, $\gamma = 120^\circ$ | ancGR2 |
| 1M2Z | P61 | 2.5 | a = b = 125.8, c = 86.0 Å, $\gamma = 120^\circ$ | hGR_1M2Z |
| 3CLD | P61 | 2.84 | a = b = 127.3, c = 77.8 Å, $\gamma = 120^\circ$ | hGR_3CLD |
| 3E7C | P61 | 2.15 | a = b = 126.6, c = 79.0 Å, $\gamma = 120^\circ$ | hGR_3CLD |
| 3K22 | P61 | 2.1 | a = b = 127.6, c = 78.2 Å, $\gamma = 120^\circ$ | hGR_3CLD |
| 4P6X | P61 | 2.5 | a = b = 220.8, c = 74.2 Å, $\gamma = 120^\circ$ | hGR_4P6X |
| 5UC1 | P61 | 2.35 | a = b = 48.4, c = 385.8 Å, $\gamma = 120^\circ$ | HetgaGRβ |
| This work | P61 | 2.7 | a = b = 190.6, c = 137.9 Å, $\gamma = 120^\circ$ | ancGR2 |
| 3BQD | P62 | 2.5 | a = b = 93.8, c = 130.0 Å, $\gamma = 120^\circ$ | hGR_1M2Z |
| 3MNE | P65 | 1.96 | a = b = 71.8, c = 128.6 Å, $\gamma = 120^\circ$ | mGR |
| 3MNP | P65 | 1.5 | a = b = 72.1, c = 128.8 Å, $\gamma = 120^\circ$ | mGR |
| 3MNO | P65 | 1.55 | a = b = 71.4, c = 127.9 Å, $\gamma = 120^\circ$ | mGR |
| 3H52 | P3221 | 2.8 | a = b = 99.5, c = 252.4 Å, $\gamma = 120^\circ$ | hGR_3H52 |
| 4MDD | P3221 | 2.4 | a = b = 72.5, c = 229.5 Å, $\gamma = 120^\circ$ | hGR_3H52 |
| 4CSJ | P3221 | 2.3 | a = b = 84.4, c = 105.7 Å, $\gamma = 120^\circ$ | hGR_4CSJ |
| 4UDC | P3221 | 2.5 | a = b = 84.7, c = 105.9 Å, $\gamma = 120^\circ$ | hGR_1NHZ |
| 4UDD | P3221 | 1.8 | a = b = 87.2, c = 102.9 Å, $\gamma = 120^\circ$ | hGR_4CSJ |
| 5G3J | P3221 | 2.4 | a = b = 84.3, c = 106.7 Å, $\gamma = 120^\circ$ | hGR_4CSJ |
| 5G5W | P3221 | 2.2 | a = b = 83.9, c = 105.7 Å, $\gamma = 120^\circ$ | hGR_4CSJ |
| 5NFP | P3221 | 2.1 | a = b = 84.0, c = 106.1 Å, $\gamma = 120^\circ$ | hGR_4CSJ |
| 5NFT | P3221 | 2.3 | a = b = 84.4, c = 106.1 Å, $\gamma = 120^\circ$ | hGR_5NFT |
| 6EL6 | P3221 | 2.4 | a = b = 85.0, c = 106.4 Å, $\gamma = 120^\circ$ | hGR_4CSJ |
| 6EL7 | P3221 | 2.18 | a = b = 83.8, c = 106.0 Å, $\gamma = 120^\circ$ | hGR_5NFT |
| 6EL9 | P3221 | 2.19 | a = b = 84.5, c = 105.7 Å | hGR_5NFT |
| This work | I4122 | 3.18 | a = b = 180.3, c = 169.2 Å | ancGR2 |
| 4P6W | P23 | 1.95 | a = b = c = 130.2 Å | hGR_4P6W |
| This work | I4132 | 3.36 | a = b = c = 178.05 Å | ancGR2 |

[1] Sequence not identical to ancGR1 – rather an intermediate between ancGR1 and ancGR2.
[2] His-Ser-Ser-Arg-Leu-Trp-Glu-Leu-Leu-Met-Glu-Ala-Thr. (Leucines of the LXXLL motif are underlined).
[3] Structure not refined. (R 34.5%, Rfree 36.3%, and worrisome MolProbity statistics).
[4] Only residues Asn742-Asp752 defined by electron density.

**Table S3.**
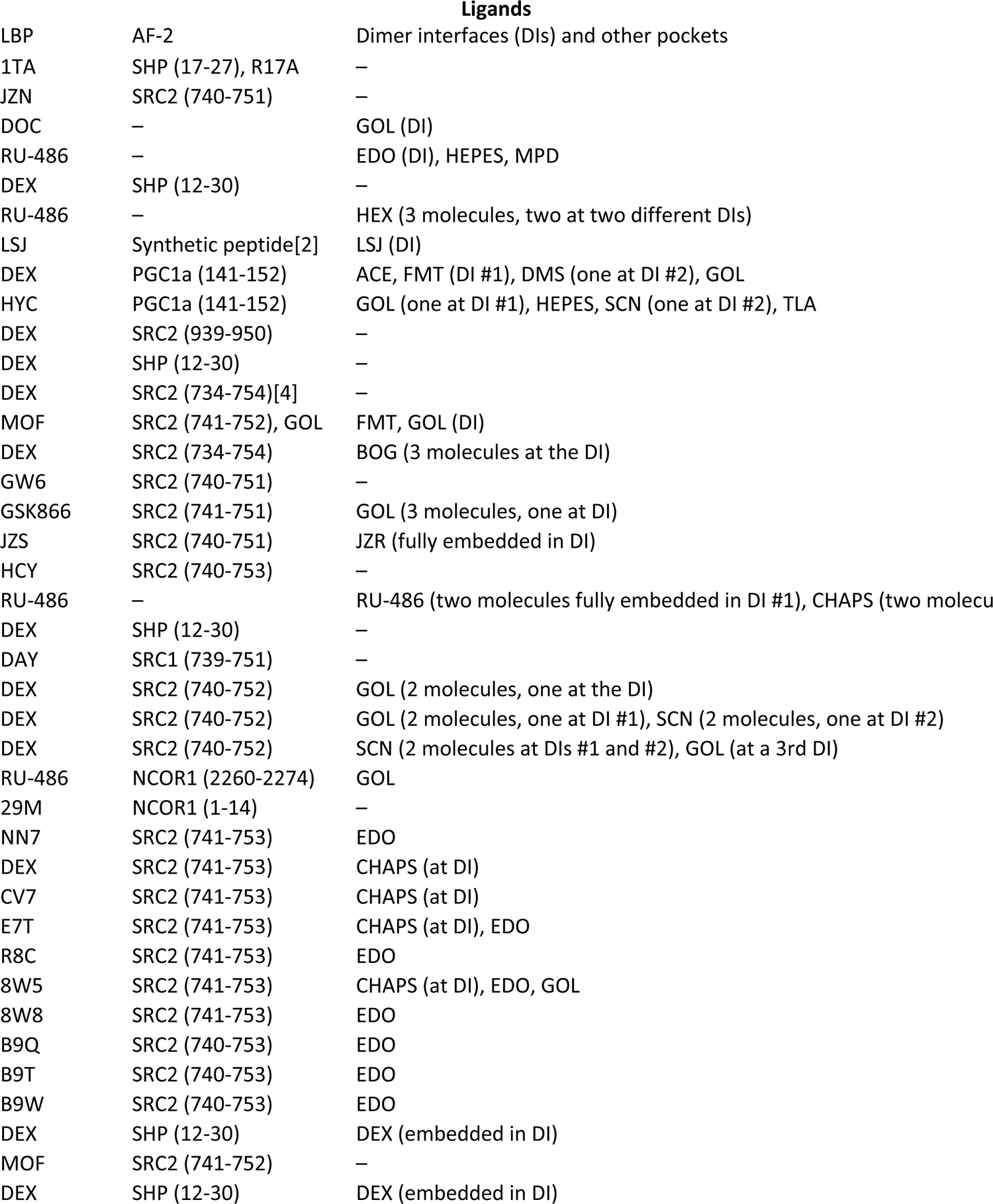
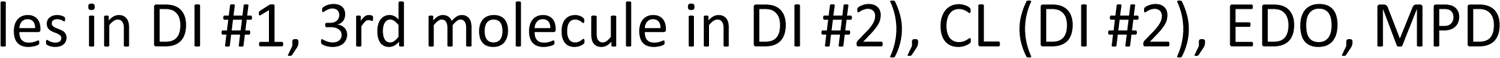
legend. Heatmap representation of the 20 topologically distinct GR-LBD homodimers. Interface residues in all pairs of GR-LBD monomers identified by PISA are marked (X).

**Table S4.**
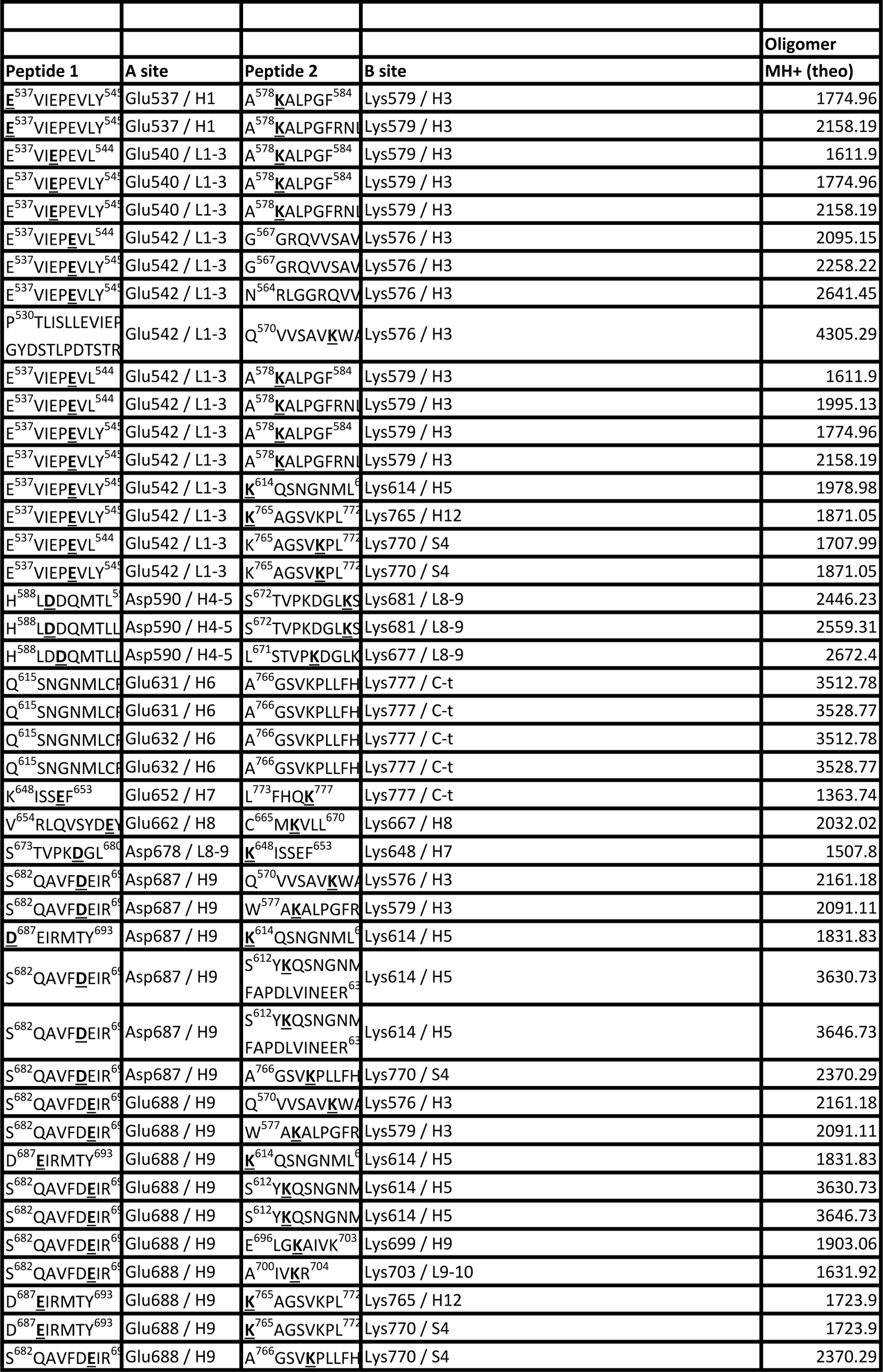
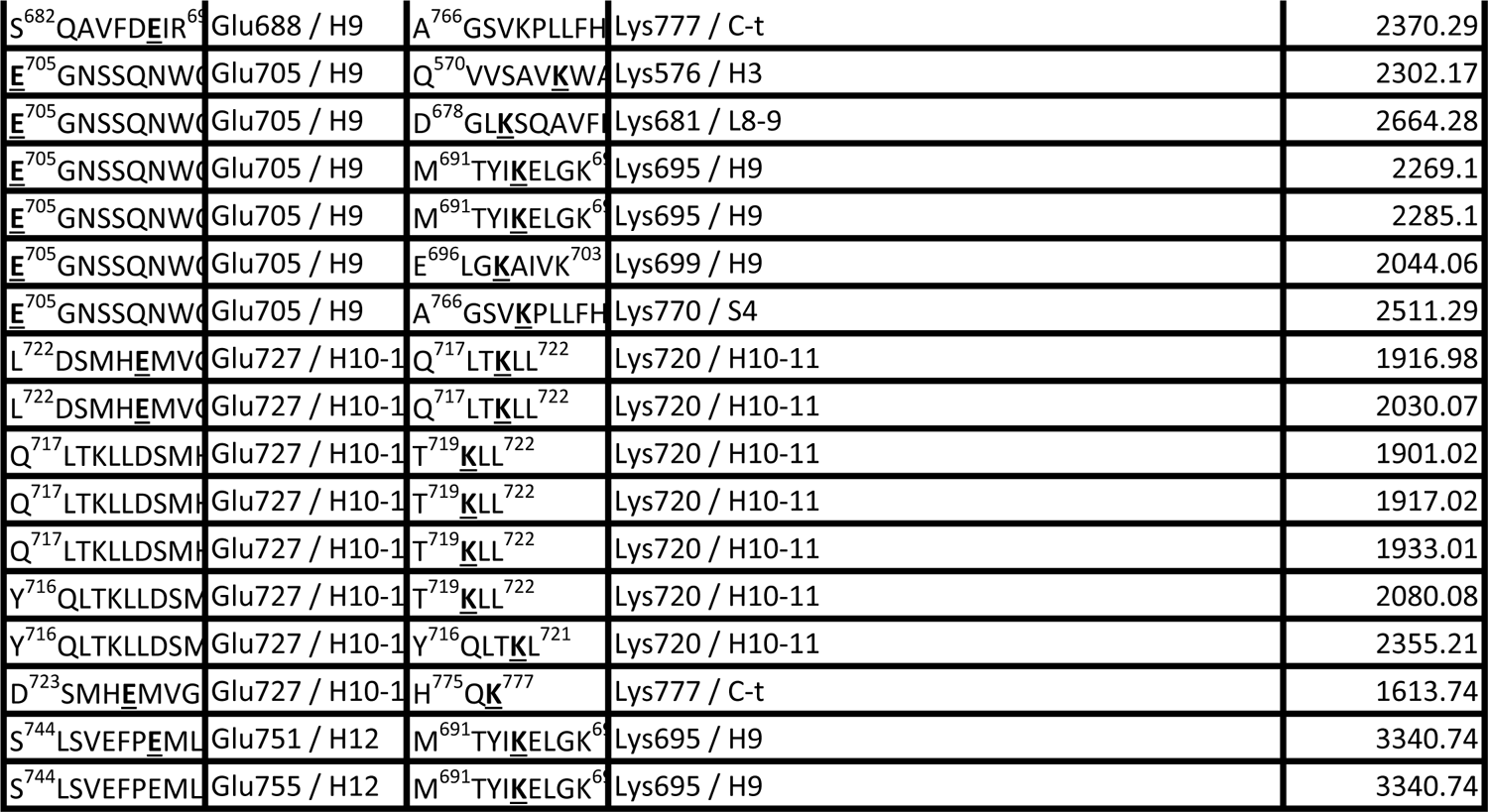
legend. Summary of EDC-crosslinked peptides of ancGR2-LBD identified by mass spectrometry.

**Table S5.** legend. MS/MS verification of EDC-mediated crosslink between residues Glu688 and Lys699 The masses of generated a, b and y ions are indicated.

| Count |  | Average MH+ (exp) |
| --- | --- | --- |
| Dim | Tetra | Dimer |
| 1 |  | 1774.96228 |
| 1 |  | 2158.18928 |
| 1 |  | 1611.89828 |
| 1 |  | 1774.96228 |
| 1 |  | 2158.18828 |
| 2 |  | 2095.15378 |
| 2 |  | 2258.21778 |
| 1 |  | 2641.45228 |
| 2 |  | 4305.27978 |
| 1 |  | 1611.90028 |
| 1 |  | 1995.12728 |
|  | 1 |  |
| 2 |  | 2158.18993 |
| 1 |  | 1978.97828 |
| 1 |  | 1871.05128 |
| 1 |  | 1707.98828 |
| 2 |  | 1871.05178 |
| 3 |  | 2446.23094 |
| 2 |  | 2559.31028 |
| 1 |  | 2672.39628 |
| 5 |  | 3512.73379 |
| 1 |  | 3528.76428 |
| 2 |  | 3512.73745 |
| 1 |  | 3528.75828 |
| 1 |  | 1363.73516 |
| 1 |  | 2032.02228 |
| 1 |  | 1507.80235 |
| 2 |  | 2161.16389 |
| 1 |  | 2091.11328 |
| 1 |  | 1831.83028 |
| 2 |  | 3630.73453 |
| 2 |  | 3646.73477 |
|  | 1 |  |
| 6 | 1 | 2161.17542 |
| 1 |  | 2091.11428 |
| 2 |  | 1831.82678 |
| 3 |  | 3630.72128 |
| 3 |  | 3646.70528 |
| 11 | 2 | 1903.06413 |
| 4 |  | 1631.9221 |
| 1 |  | 1723.90264 |
|  | 1 |  |
|  | 2 |  |
| 1 |  | 2370.29628 |
| 2 |  | 2302.16954 |
|  | 1 |  |
| 9 |  | 2269.1015 |
| 9 | 2 | 2285.09668 |
| 1 |  | 2044.05828 |
| 16 |  | 2511.28502 |
| 1 |  | 1916.98128 |
| 1 |  | 2030.06528 |
| 5 |  | 1900.98772 |
| 5 | 2 | 1916.98137 |
| 5 |  | 1932.976 |
| 1 |  | 2080.04398 |
| 5 |  | 2355.17114 |
| 1 |  | 1613.74016 |
| 2 |  | 3340.74336 |
| 1 |  | 3340.74185 |

**Table S6.**
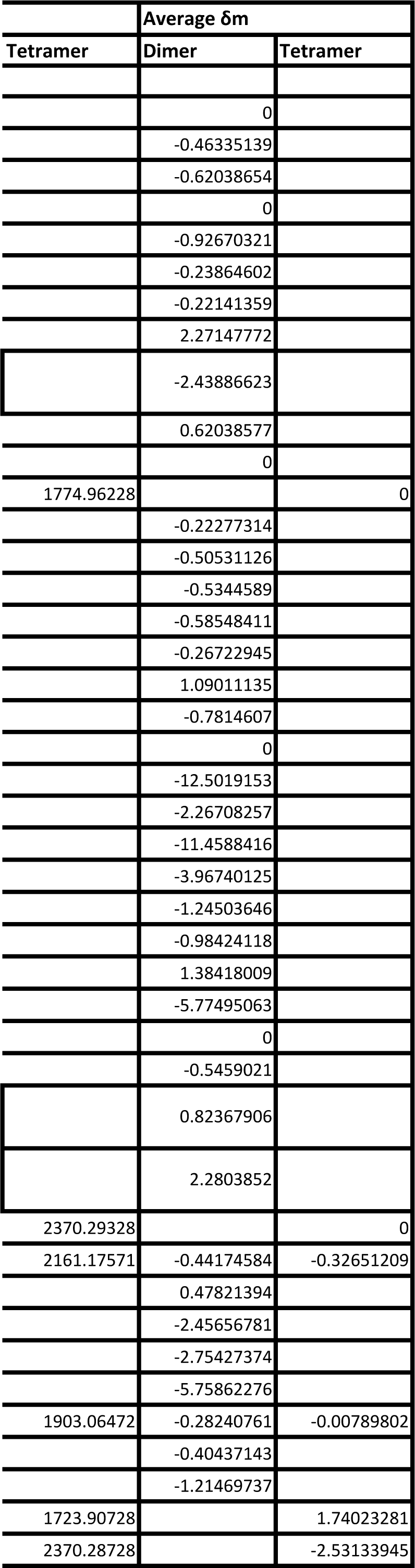
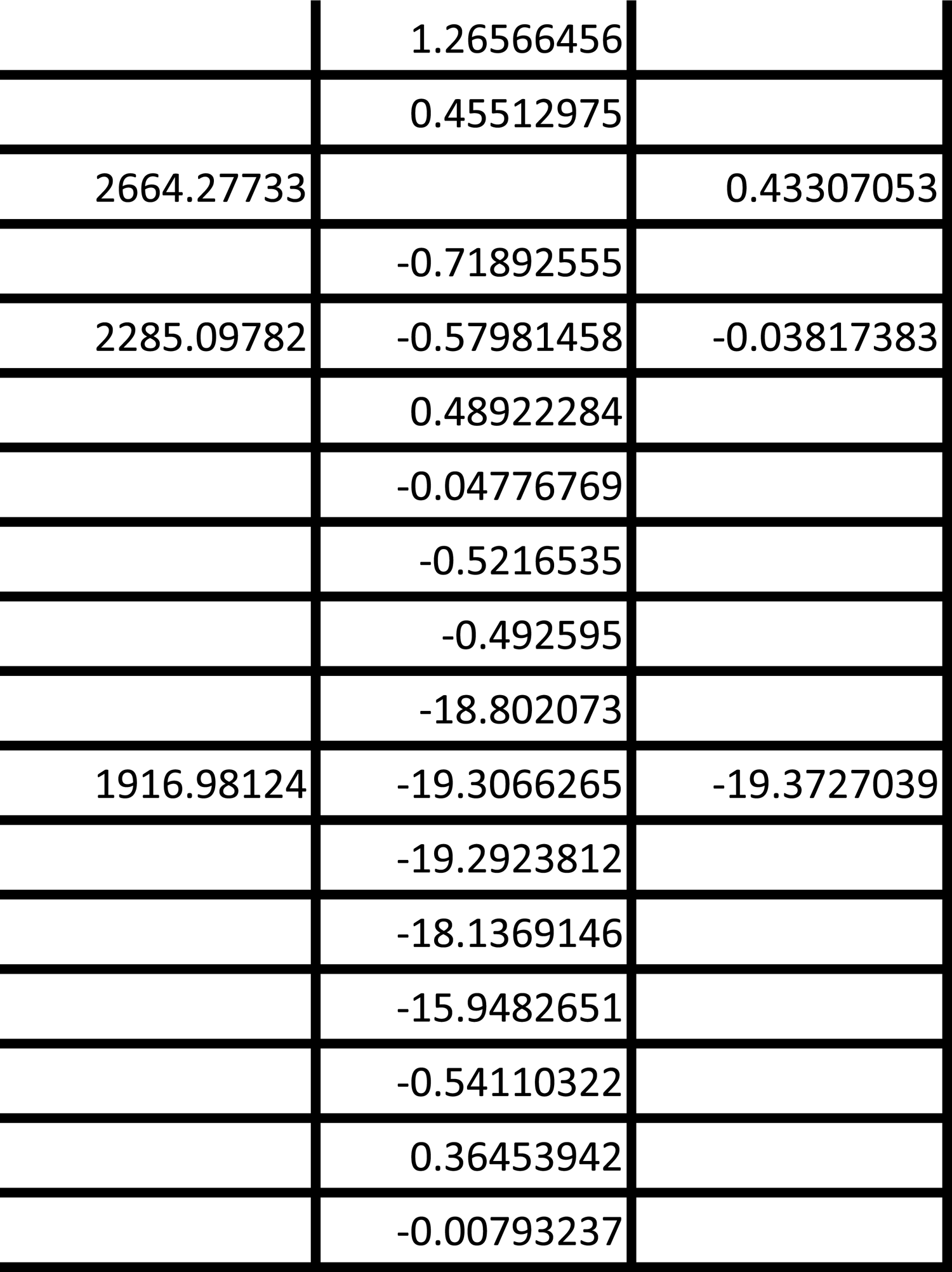
legend. Summary of DSBU-crosslinked peptides of ancGR2-LBD identified by mass spectrometry.

**Table S7.**
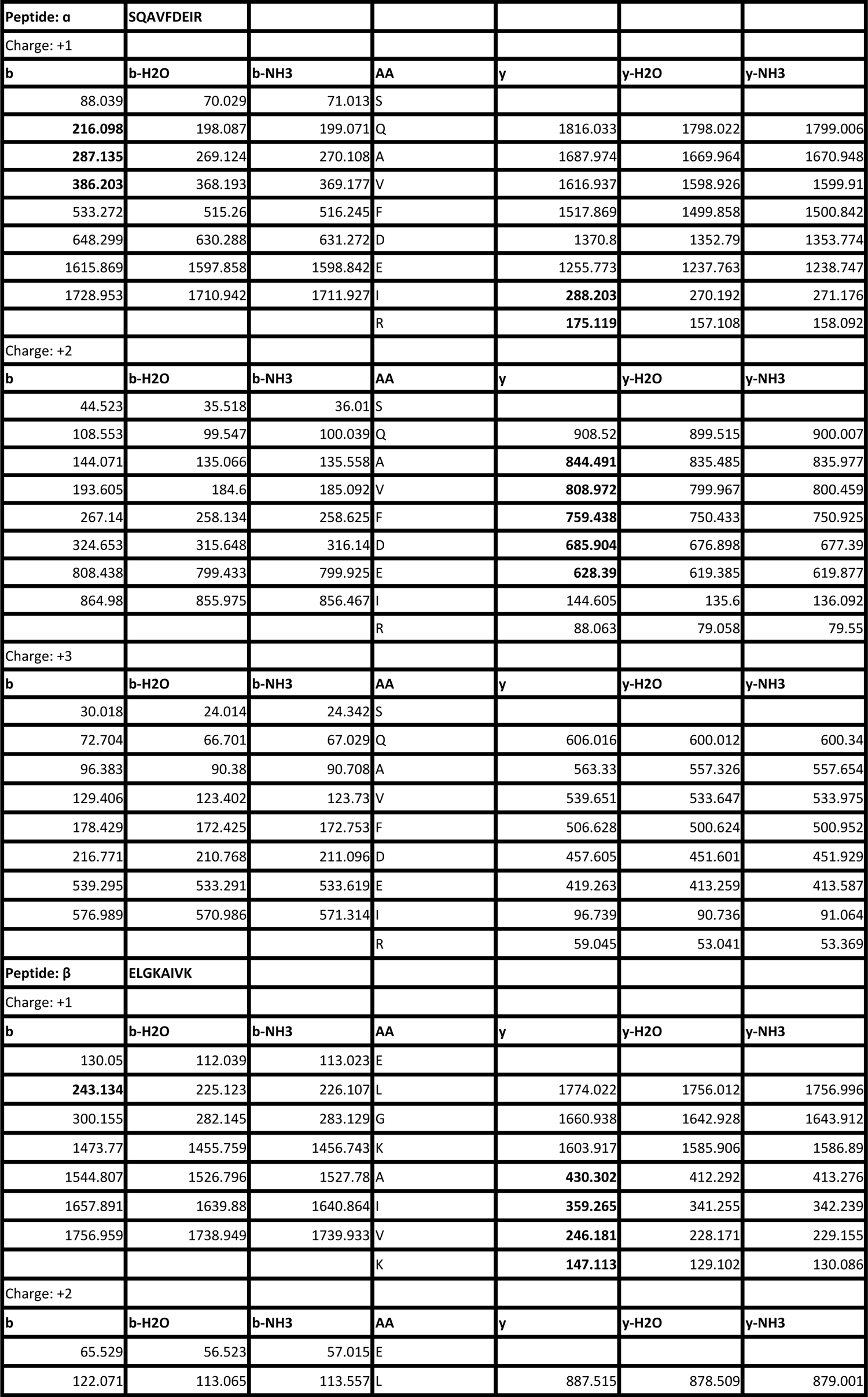
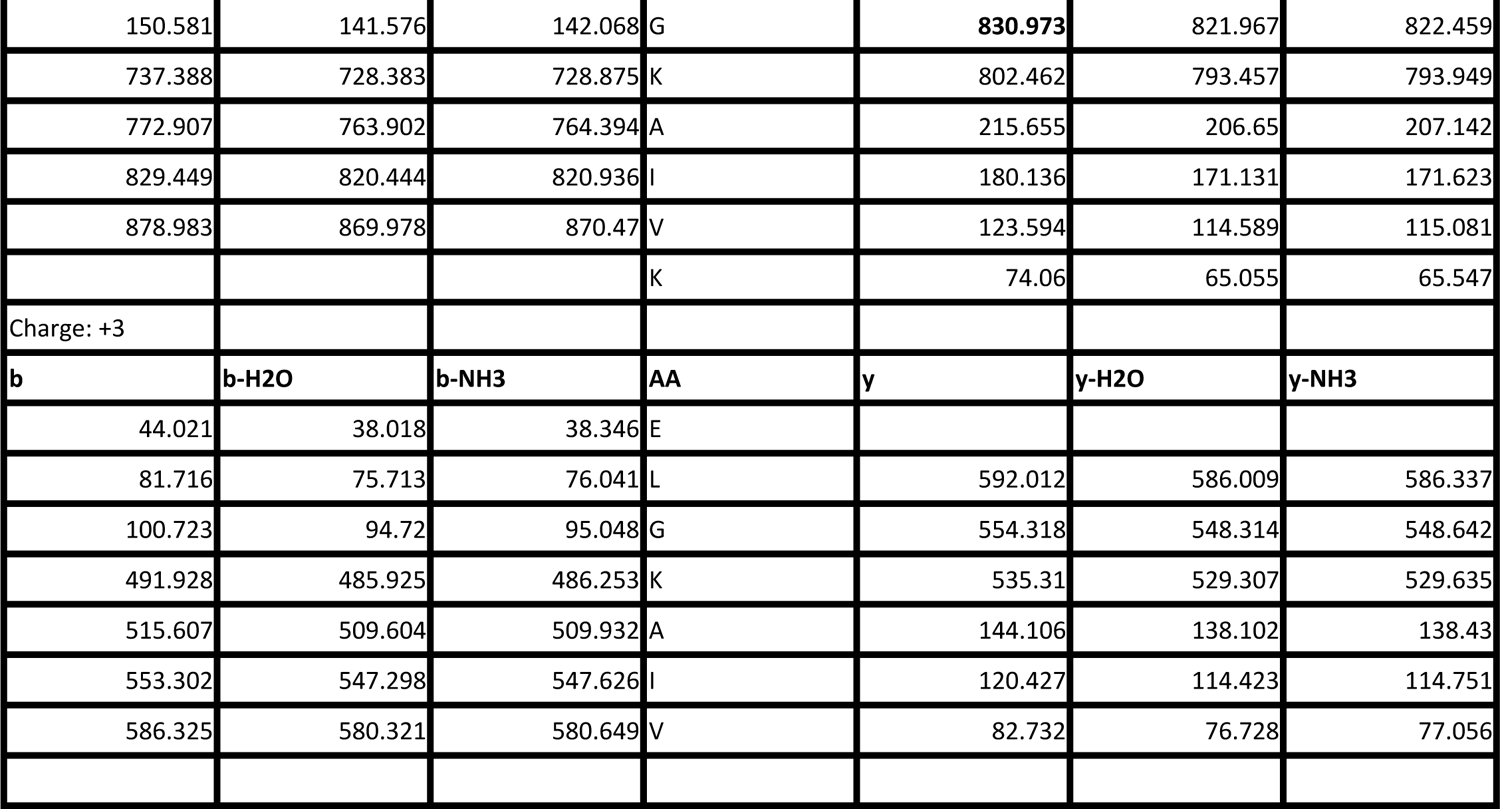
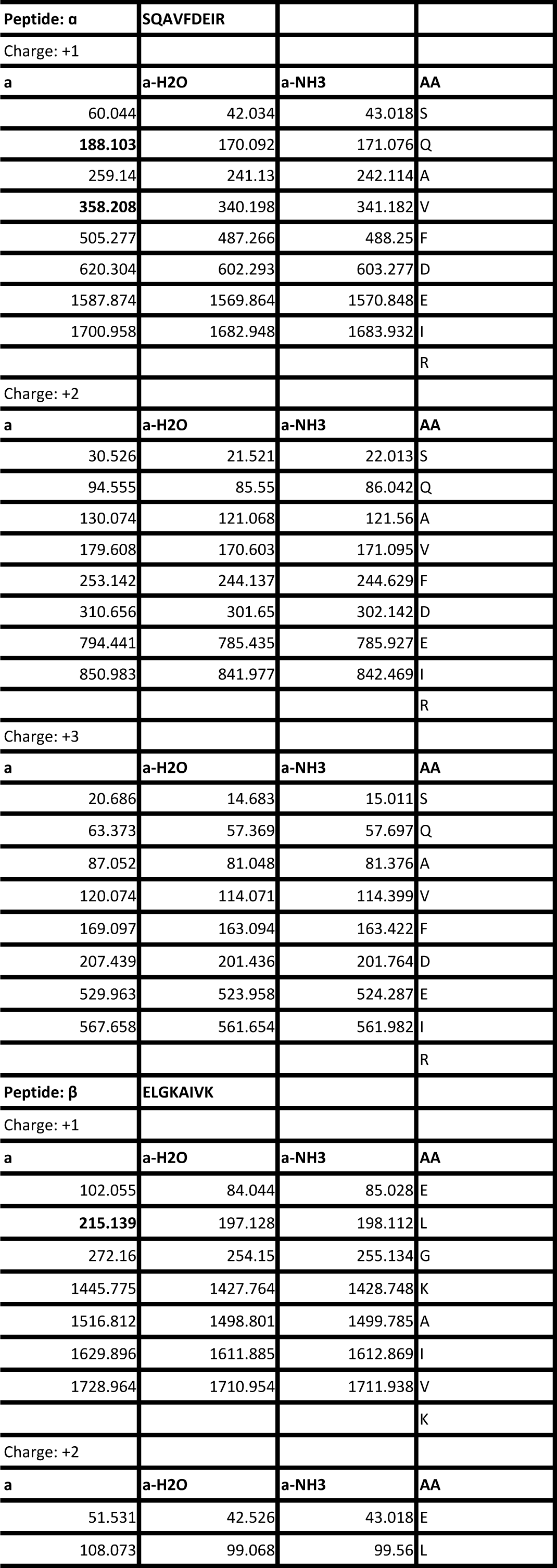
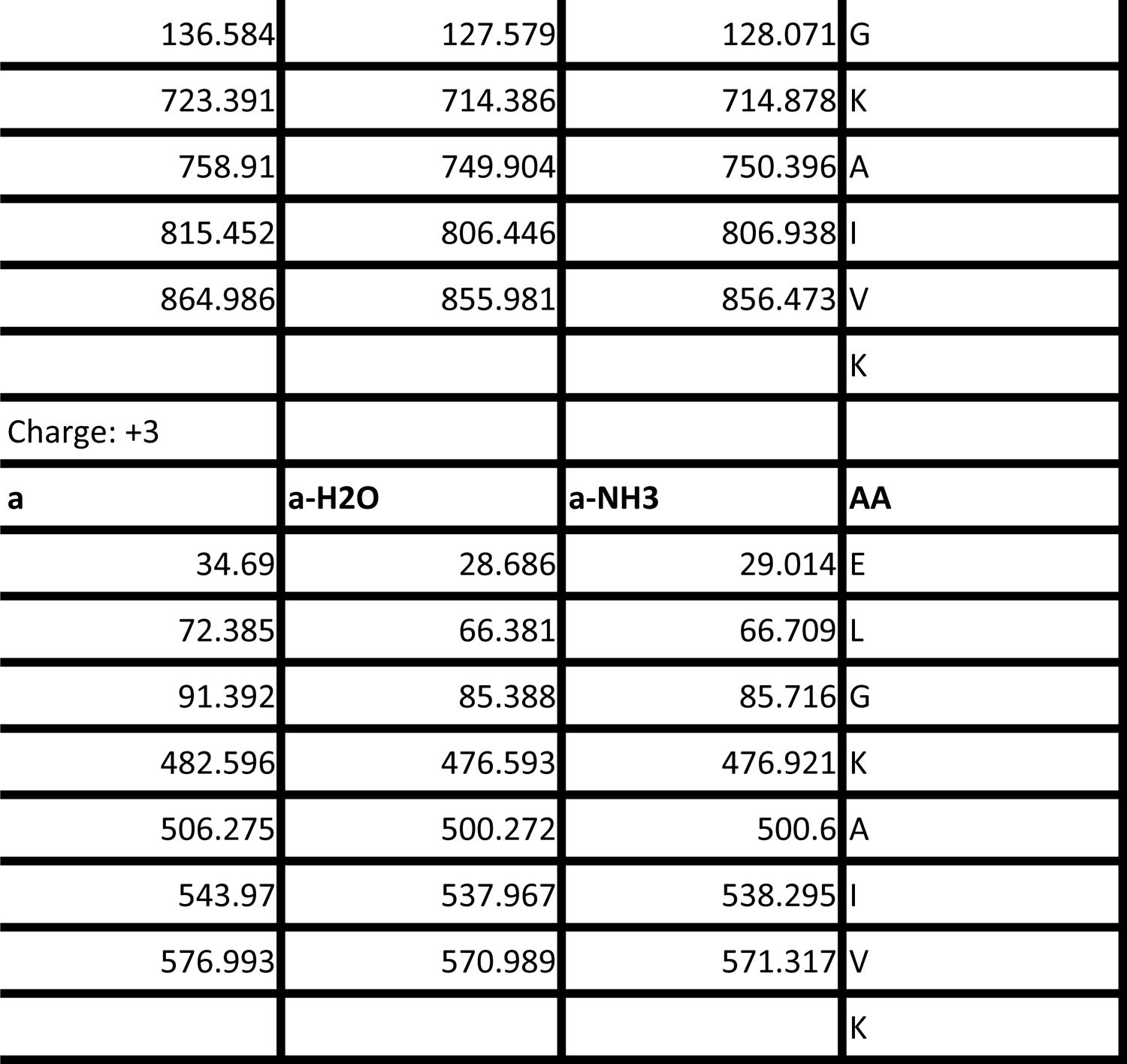
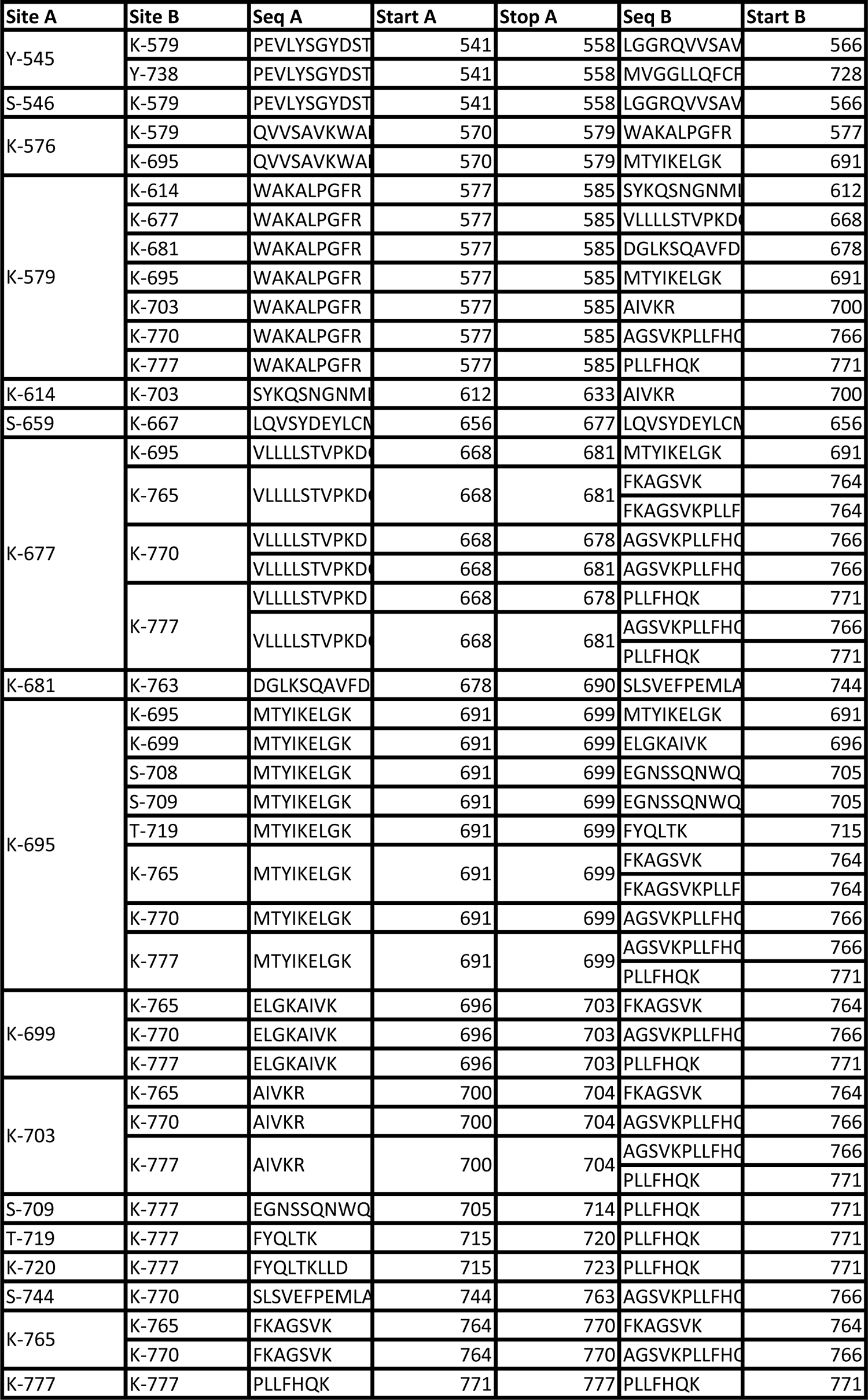
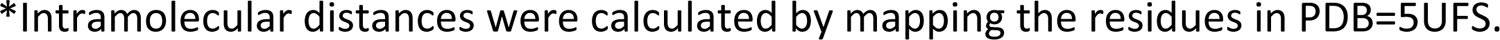
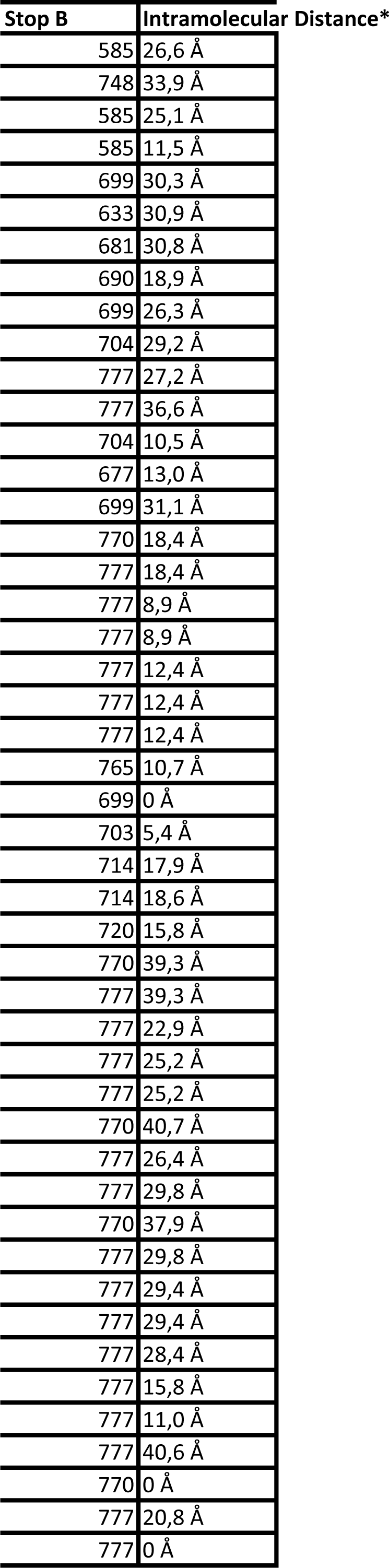
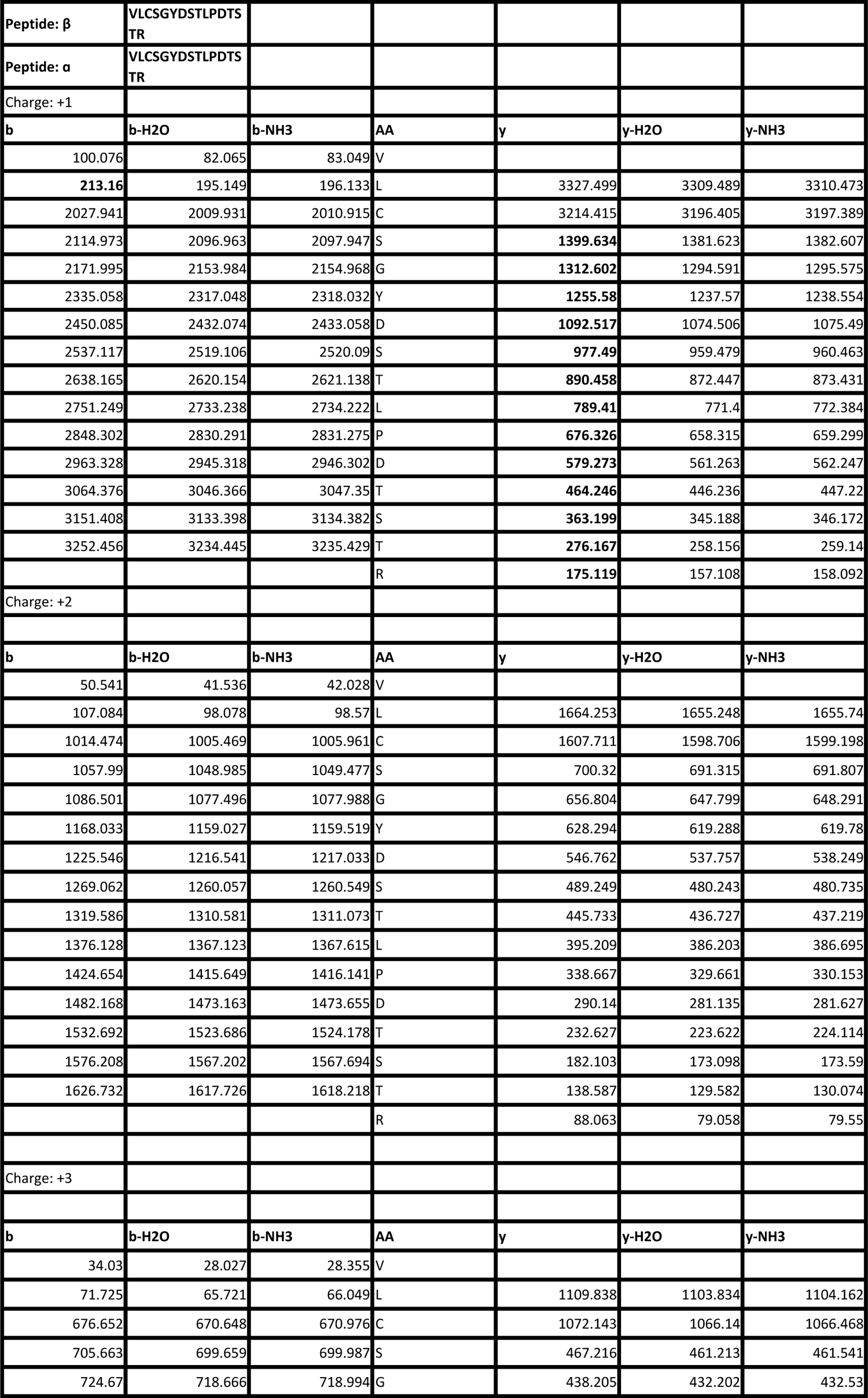
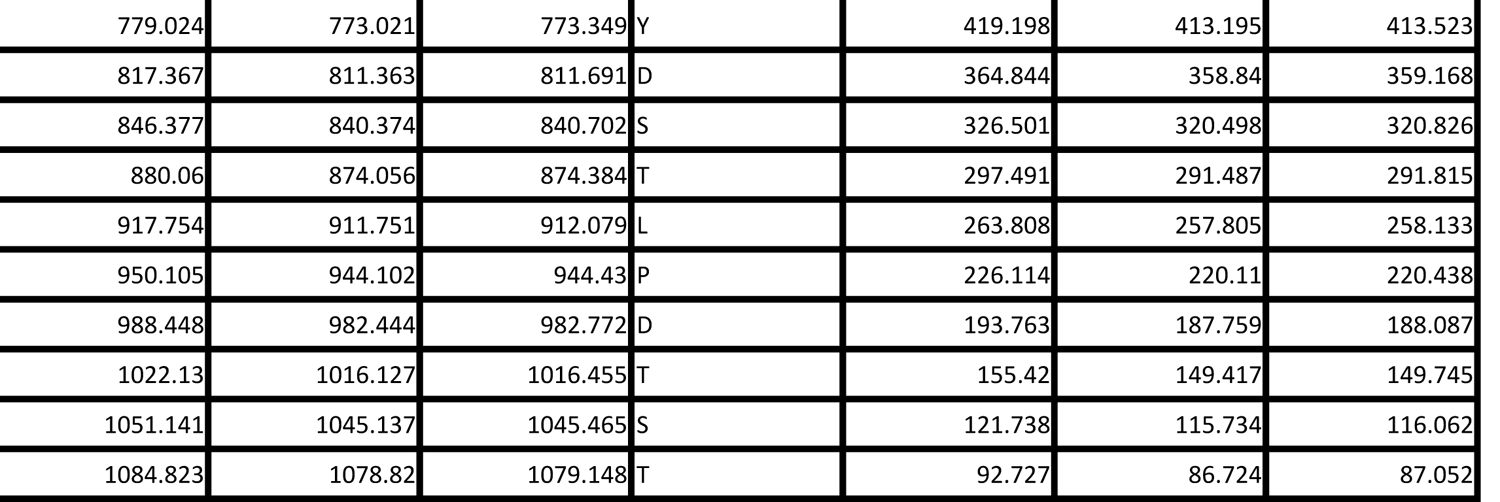
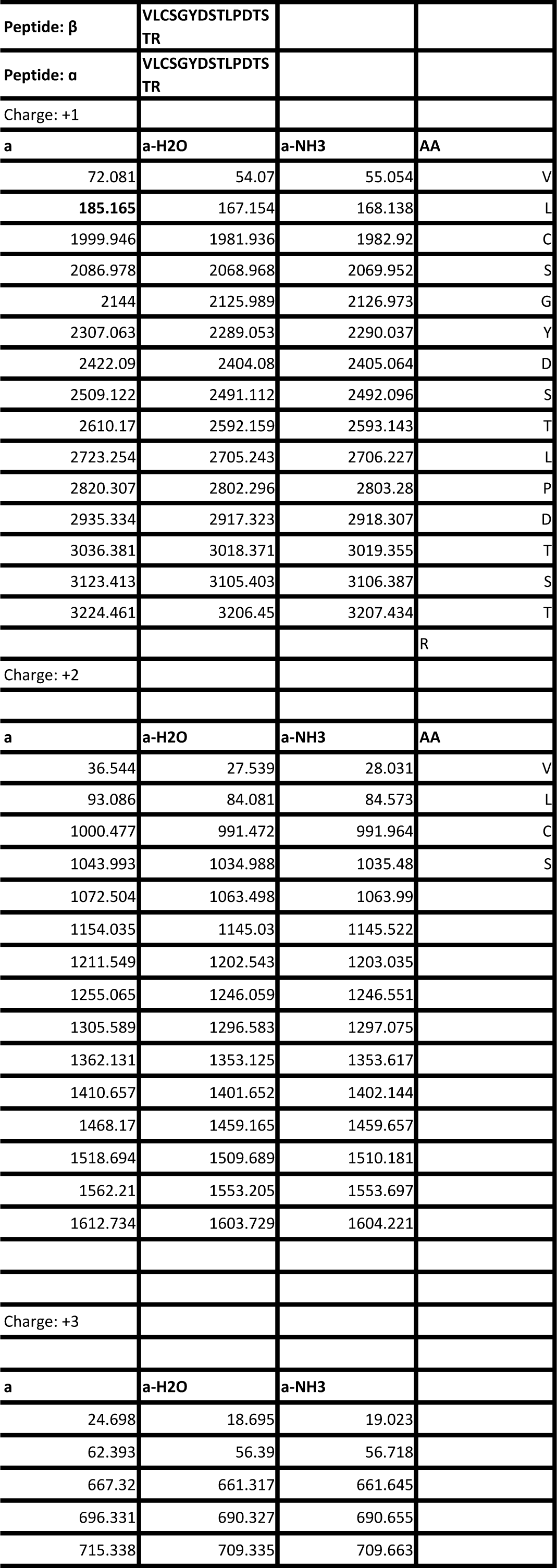
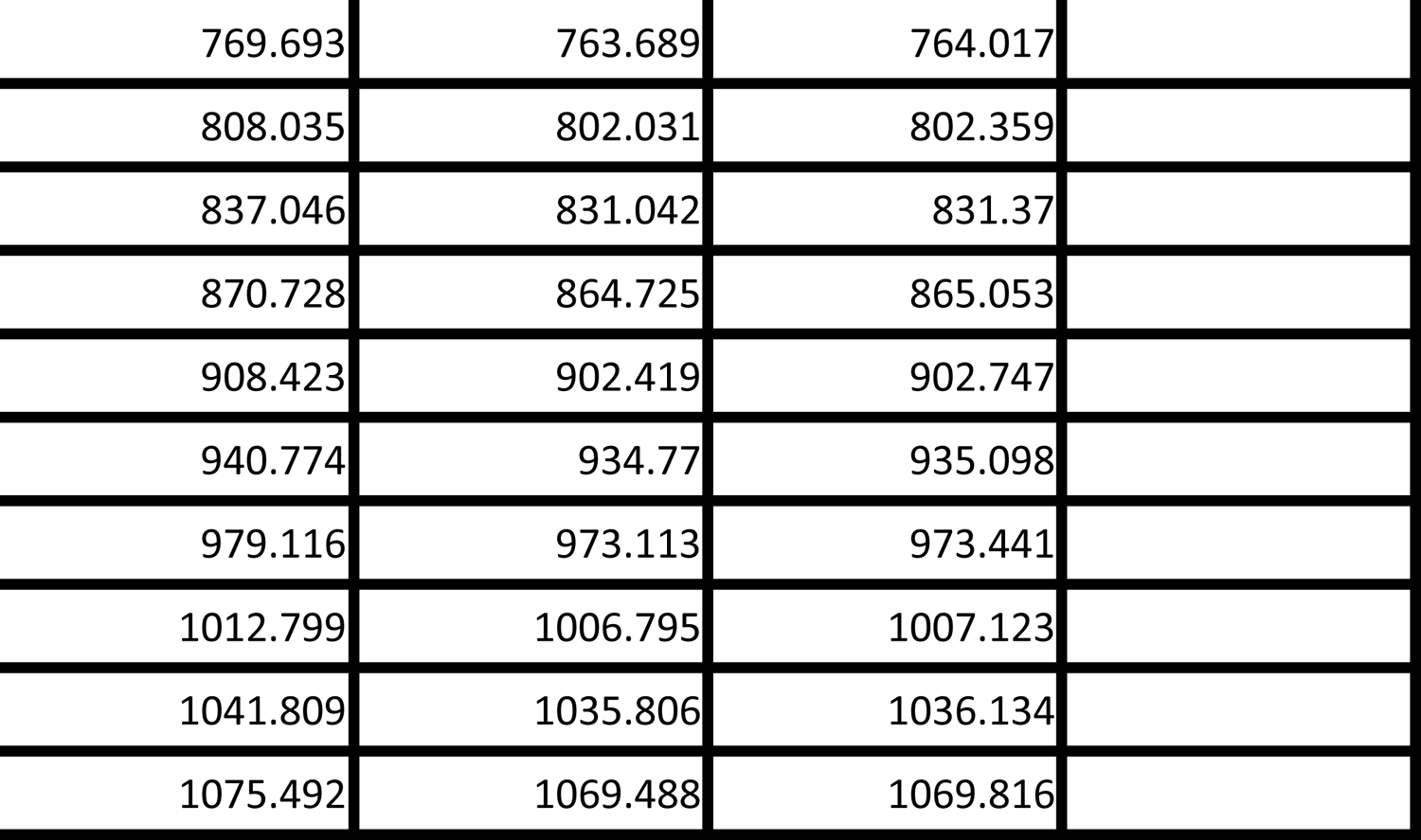
legend. MS/MS verification of disulfide bridge formation between Cys545 residues from two GR-LBD(Y545C) monomers. The masses of generated a, b and y ions are given.

## References

1. Bianchetti L, Wassmer B, Defosset A, Smertina A, Tiberti ML, Stote RH & Dejaegere A (2018) Alternative dimerization interfaces in the glucocorticoid receptor-α ligand binding domain. Biochimica et Biophysica Acta - General Subjects 1862: 1810–1825

2. Biggadike K, Bledsoe RK, Coe DM, Cooper TWJ, House D, Iannone MA, Macdonald SJF, Madauss KP, McLay IM, Shipley TJ, et al (2009) Design and x-ray crystal structures of high-potency nonsteroidal glucocorticoid agonists exploiting a novel binding site on the receptor. Proceedings of the National Academy of Sciences of the United States of America 106: 18114– 18119

3. Bledsoe RK, Montana VG, Stanley TB, Delves CJ, Apolito CJ, McKee DD, Consler TG, Parks DJ, Stewart EL, Willson TM, et al (2002) Crystal structure of the glucocorticoid receptor ligand binding domain reveals a novel mode of receptor dimerization and coactivator recognition. Cell 110: 93–105

4. Bogan AA & Thorn KS (1998) Anatomy of hot spots in protein interfaces. Journal of Molecular Biology 280: 1–9

5. De Bosscher K, Desmet SJ, Clarisse D, Estébanez-Perpiña E & Brunsveld L (2020) Nuclear receptor crosstalk — defining the mechanisms for therapeutic innovation. Nature Reviews Endocrinology 16: 363–377 doi:10.1038/s41574-020-0349-5

6. Busada JT & Cidlowski JA (2017) Mechanisms of Glucocorticoid Action During Development. In Current Topics in Developmental Biology pp 147–170. Academic Press Inc.

7. Cain DW & Cidlowski JA (2017) Immune regulation by glucocorticoids. Nature Reviews Immunology 17: 233–247 doi:10.1038/nri.2017.1

8. Canutescu AA, Shelenkov AA & Dunbrack RL (2003) A graph-theory algorithm for rapid protein side-chain prediction. Protein Science 12: 2001–2014

9. Carroll SM, Ortlund EA & Thornton JW (2011) Mechanisms for the Evolution of a Derived Function in the Ancestral Glucocorticoid Receptor. PLoS Genetics 7: e1002117

10. Carson MW, Luz JG, Suen C, Montrose C, Zink R, Ruan X, Cheng C, Cole H, Adrian MD, Kohlman DT, et al (2014) Glucocorticoid receptor modulators informed by crystallography lead to a new rationale for receptor selectivity, function, and implications for structure-based design. Journal of Medicinal Chemistry 57: 849–860

11. Chandra V, Wu D, Li S, Potluri N, Kim Y & Rastinejad F (2017) The quaternary architecture of RARβ–RXRα heterodimer facilitates domain–domain signal transmission. Nature Communications 8: 868

12. Chen R, Li L & Weng Z (2003) ZDOCK: An initial-stage protein-docking algorithm. *Proteins: Structure*, Function and Genetics 52: 80–87

13. Cheng TMK, Blundell TL & Fernandez-Recio J (2007) PyDock: Electrostatics and desolvation for effective scoring of rigid-body protein-protein docking. *Proteins: Structure*, Function and Genetics 68: 503–515

14. Chrousos GP, Loriaux DL, Tomita M, Brandon DD, Renquist D, Albertson B & Lipsett MB (1986) The new world primates as animal models of glucocorticoid resistance. Advances in experimental medicine and biology 196: 129–144 doi:10.1007/978-1-4684-5101-6_9

15. Clark AR & Belvisi MG (2012) Maps and legends: The quest for dissociated ligands of the glucocorticoid receptor. Pharmacology and Therapeutics 134: 54–67 doi:10.1016/j.pharmthera.2011.12.004

16. Conway-Campbell BL, Pooley JR, Hager GL & Lightman SL (2012) Molecular dynamics of ultradian glucocorticoid receptor action. Molecular and Cellular Endocrinology 348: 383–393 doi:10.1016/j.mce.2011.08.014

17. Dendoncker K, Timmermans S, Vandewalle J, Eggermont M, Lempiäinen J, Van Hamme E, Dewaele S, Vandevyver S, Ballegeer M, Souffriau J, et al (2019) TNF-α inhibits glucocorticoid receptor-induced gene expression by reshaping the GR nuclear cofactor profile. Proceedings of the National Academy of Sciences of the United States of America 116: 12942–12951

18. Digman MA, Dalal R, Horwitz AF & Gratton E (2008) Mapping the number of molecules and brightness in the laser scanning microscope. Biophysical Journal 94: 2320–2332

19. Escoter-Torres L, Greulich F, Quagliarini F, Wierer M & Uhlenhaut NH (2020) Anti-inflammatory functions of the glucocorticoid receptor require DNA binding. Nucleic Acids Research 48: 8393–8407

20. Estebanez-Perpina E, Arnold LA, Nguyen P, Rodrigues ED, Mar E, Bateman R, Pallai P, Shokat KM, Baxter JD, Guy RK, et al (2007) A surface on the androgen receptor that allosterically regulates coactivator binding. Proceedings of the National Academy of Sciences 104: 16074–16079

21. Evans RM & Mangelsdorf DJ (2014) Nuclear receptors, RXR, and the big bang. Cell 157: 255–266

22. Fernandez-Recio J, Totrov M, Skorodumov C & Abagyan R (2005) Optimal docking area: A new method for predicting protein-protein interaction sites. *Proteins: Structure*, Function and Genetics 58: 134–143

23. Franco LM, Gadkari M, Howe KN, Sun J, Kardava L, Kumar P, Kumari S, Hu Z, Fraser IDC, Moir S, et al (2019) Immune regulation by glucocorticoids can be linked to cell type–dependent transcriptional responses. Journal of Experimental Medicine 216: 384–406

24. Frank F, Okafor CD & Ortlund EA (2018) The first crystal structure of a DNA-free nuclear receptor DNA binding domain sheds light on DNA-driven allostery in the glucocorticoid receptor. Scientific Reports 8

25. Fuentes-Prior P, Rojas A, Hagler AT & Estébanez-Perpiñá E (2019) Diversity of Quaternary Structures Regulates Nuclear Receptor Activities. Trends in Biochemical Sciences 44: 2–6 doi:10.1016/j.tibs.2018.09.005

26. Gabb HA, Jackson RM & Sternberg MJE (1997) Modelling protein docking using shape complementarity, electrostatics and biochemical information. Journal of Molecular Biology 272: 106–120

27. Gallastegui N, Mackinnon JAG, Fletterick RJ & Estébanez-Perpiñá E (2015) Advances in our structural understanding of orphan nuclear receptors. Trends in Biochemical Sciences 40: 25–35

28. Gampe RT, Montana VG, Lambert MH, Wisely GB, Milburn M V & Xu HE (2000) Structural basis for autorepression of retinoid X receptor by tetramer formation and the AF-2 helix. Genes & development 14: 2229–41

29. Garcia DA, Johnson TA, Presman DM, Fettweis G, Wagh K, Rinaldi L, Stavreva DA, Paakinaho V, Jensen RAM, Mandrup S, et al (2021) An intrinsically disordered region-mediated confinement state contributes to the dynamics and function of transcription factors. Molecular Cell 81: 1484–1498.e6

30. Grosdidier S & Fernández-Recio J (2008) Identification of hot-spot residues in protein-protein interactions by computational docking. BMC Bioinformatics 9

31. Halabi N, Rivoire O, Leibler S & Ranganathan R (2009) Protein Sectors: Evolutionary Units of Three-Dimensional Structure. Cell 138: 774–786

32. Härd T, Kellenbach E, Boelens R, Maler BA, Dahlman K, Freedman LP, Carlstedt-Duke J, Yamamoto KR, Gustafsson JÅ & Kaptein R (1990) Solution structure of the glucocorticoid receptor DNA-binding domain. Science 249: 157–160

33. He Y, Yi W, Suino-Powell K, Zhou XE, Tolbert WD, Tang X, Yang J, Yang H, Shi J, Hou L, et al (2014) Structures and mechanism for the design of highly potent glucocorticoids. Cell Research 24: 713–726

34. Housley PR, Sanchez ER, Danielsen M, Ringold GM & Pratt WB (1990) Evidence that the conserved region in the steroid binding domain of the glucocorticoid receptor is required for both optimal binding of hsp90 and protection from proteolytic cleavage. A two-site model for hsp90 binding to the steroid binding domain. The Journal of biological chemistry 265: 12778–81

35. Hudson WH, Youn C & Ortlund EA (2013) The structural basis of direct glucocorticoid-mediated transrepression. Nature Structural and Molecular Biology 20: 53–58

36. Hurley DM, Accili D, Stratakis CA, Karl M, Vamvakopoulos N, Rorer E, Constantine K, Taylor SI & Chrousos GP (1991) Point mutation causing a single amino acid substitution in the hormone binding domain of the glucocorticoid receptor in familial glucocorticoid resistance. Journal of Clinical Investigation 87: 680–686

37. Hurt DE, Suzuki S, Mayama T, Charmandari E & Kino T (2016) Structural analysis on the pathologic mutant glucocorticoid receptor ligand-binding domains. Molecular Endocrinology 30: 173–188

38. Jehle K, Cato L, Neeb A, Muhle-Goll C, Jung N, Smith EW, Buzon V, Carbó LR, Estébanez-Perpiñá E, Schmitz K, et al (2014) Coregulator control of androgen receptor action by a novel nuclear receptor-binding motif. Journal of Biological Chemistry 289: 8839–8851

39. Jiménez-Panizo A, Pérez P, Rojas AM, Fuentes-Prior P & Estébanez-Perpiñá E (2019) Non-canonical dimerization of the androgen receptor and other nuclear receptors: Implications for human disease. Endocrine-Related Cancer 26: R479–R497 doi:10.1530/ERC-19-0132

40. Johnson TA, Paakinaho V, Kim S, Hager GL & Presman DM (2021) Genome-wide binding potential and regulatory activity of the glucocorticoid receptor’s monomeric and dimeric forms. Nature Communications 12

41. Kauppi B, Jakob C, Färnegårdh M, Yang J, Ahola H, Alarcon M, Calles K, Engström O, Harlan J, Muchmore S, et al (2003) The three-dimensional structures of antagonistic and agonistic forms of the glucocorticoid receptor ligand-binding domain: RU-486 induces a transconformation that leads to active antagonism. Journal of Biological Chemistry 278: 22748–22754

42. Liu X, Wang Y & Ortlund EA (2019) First high-resolution crystal structures of the glucocorticoid receptor ligand-binding domain–peroxisome proliferator-activated γ coactivator 1-α complex with endogenous and synthetic glucocorticoids. Molecular Pharmacology 96: 408–417

43. Lockless SW & Ranganathan R (1999) Evolutionarily conserved pathways of energetic connectivity in protein families. Science 286: 295–299

44. Louw A (2019) GR dimerization and the impact of gr dimerization on gr protein stability and half-life. Frontiers in Immunology 10 doi:10.3389/fimmu.2019.01693

45. Luisi BF, Xu WX, Otwinowski Z, Freedman LP, Yamamoto KR & Sigler PB (1991) Crystallographic analysis of the interaction of the glucocorticoid receptor with DNA. Nature 352: 497–505

46. McNally JC, Müller WG, Walker D, Wolford R & Hager GL (2000) The glucocorticoid receptor: Rapid exchange with regulatory sites in living cells. Science 287: 1262–1265

47. Meijsing SH, Pufall MA, So AY, Bates DL, Chen L & Yamamoto KR (2009) DNA binding site sequence directs glucocorticoid receptor structure and activity. *Science (New York*, NY) 324: 407–10

48. Mikuni S, Tamura M & Kinjo M (2007) Analysis of intranuclear binding process of glucocorticoid receptor using fluorescence correlation spectroscopy. FEBS Letters 581: 389–393

49. Min J, Perera L, Krahn JM, Jewell CM, Moon AF, Cidlowski JA & Pedersen LC (2018) Probing Dominant Negative Behavior of Glucocorticoid Receptor β through a Hybrid Structural and Biochemical Approach. Molecular and Cellular Biology 38: e00453–17

50. Nadal M, Prekovic S, Gallastegui N, Helsen C, Abella M, Zielinska K, Gay M, Vilaseca M, Taulès M, Houtsmuller AB, et al (2017) Structure of the homodimeric androgen receptor ligand-binding domain. Nature Communications 8

51. Nicolaides NC & Charmandari E (2019) Glucocorticoid Resistance. Experientia supplementum (2012) 111: 85–102 doi:10.1007/978-3-030-25905-1_6

52. Noddings CM, Wang RYR & Agard DA (2020) GR chaperone cycle mechanism revealed by cryo-EM: Reactivation of GR by the GR:Hsp90:p23 client-maturation complex. bioRxiv: 2020.09.12.294975 doi:10.1101/2020.09.12.294975

53. Oh KS, Patel H, Gottschalk RA, Lee WS, Baek S, Fraser IDC, Hager GL & Sung MH (2017) Anti-Inflammatory Chromatinscape Suggests Alternative Mechanisms of Glucocorticoid Receptor Action. Immunity 47: 298–309.e5

54. Ortlund EA, Bridgham JT, Redinbo MR & Thornton JW (2007) Crystal structure of an ancient protein: evolution by conformational epistasis. *Science (New York*, NY) 317: 1544–8

55. Paakinaho V, Johnson TA, Presman DM & Hager GL (2019) Glucocorticoid receptor quaternary structure drives chromatin occupancy and transcriptional outcome. Genome Research 29: 1223–1234

56. Pazos F & Bang J-W (2008) Computational Prediction of Functionally Important Regions in Proteins. Current Bioinformatics 1: 15–23

57. Pfaff SJ & Fletterick RJ (2010) Hormone binding and co-regulator binding to the glucocorticoid receptor are allosterically coupled. Journal of Biological Chemistry 285: 15256–15267

58. Presman DM, Ball DA, Paakinaho V, Grimm JB, Lavis LD, Karpova TS & Hager GL (2017) Quantifying transcription factor binding dynamics at the single-molecule level in live cells. Methods 123: 76–88

59. Presman DM, Ganguly S, Schiltz RL, Johnson TA, Karpova TS & Hager GL (2016) DNA binding triggers tetramerization of the glucocorticoid receptor in live cells. Proceedings of the National Academy of Sciences 113: 8236– 8241

60. Presman DM, Ogara MF, Stortz M, Alvarez LD, Pooley JR, Schiltz RL, Grøntved L, Johnson TA, Mittelstadt PR, Ashwell JD, et al (2014) Live Cell Imaging Unveils Multiple Domain Requirements for In Vivo Dimerization of the Glucocorticoid Receptor. PLoS Biology 12

61. Rivoire O, Reynolds KA & Ranganathan R (2016) Evolution-Based Functional Decomposition of Proteins. PLoS Computational Biology 12

62. Rogatsky I, Wang JC, Derynck MK, Nonaka DF, Khodabakhsh DB, Haqq CM, Darimont BD, Garabedian MJ & Yamamoto KR (2003) Target-specific utilization of transcriptional regulatory surfaces by the glucocorticoid receptor. Proceedings of the National Academy of Sciences of the United States of America 100: 13845–13850

63. Rojas AM, Fuentes G, Rausell A & Valencia A (2012) The Ras protein superfamily: Evolutionary tree and role of conserved amino acids. Journal of Cell Biology 196: 189–201 doi:10.1083/jcb.201103008

64. Schäcke H, Berger M, Rehwinkel H & Asadullah K (2007) Selective glucocorticoid receptor agonists (SEGRAs): Novel ligands with an improved therapeutic index. Molecular and Cellular Endocrinology 275: 109–117

65. Schoch GA, D’Arcy B, Stihle M, Burger D, Bär D, Benz J, Thoma R & Ruf A (2010) Molecular Switch in the Glucocorticoid Receptor: Active and Passive Antagonist Conformations. Journal of Molecular Biology 395: 568–577

66. Sevilla LM, Bayo P, Latorre V, Sanchis A & Pérez P (2010) Glucocorticoid receptor regulates overlapping and differential gene subsets in developing and adult skin. Molecular Endocrinology 24: 2166–2178

67. Stortz M, Pecci A, Presman DM & Levi V (2020) Unraveling the molecular interactions involved in phase separation of glucocorticoid receptor. BMC Biology 18: 1–20

68. Vandevyver S, Dejager L & Libert C (2012) On the Trail of the Glucocorticoid Receptor: Into the Nucleus and Back. Traffic 13: 364–374 doi:10.1111/j.1600-0854.2011.01288.x

69. Weikum ER, Knuesel MT, Ortlund EA & Yamamoto KR (2017a) Glucocorticoid receptor control of transcription: Precision and plasticity via allostery. Nature Reviews Molecular Cell Biology 18: 159–174 doi:10.1038/nrm.2016.152

70. Weikum ER, Okafor CD, D’Agostino EH, Colucci JK & Ortlund EA (2017b) Structural analysis of the glucocorticoid receptor ligand-binding domain in complex with triamcinolone acetonide and a fragment of the atypical coregulator, small heterodimer partner. Molecular Pharmacology 92: 12–21

71. Weikum ER, de Vera IMS, Nwachukwu JC, Hudson WH, Nettles KW, Kojetin DJ & Ortlund EA (2017c) Tethering not required: The glucocorticoid receptor binds directly to activator protein-1 recognition motifs to repress inflammatory genes. Nucleic Acids Research 45: 8596–8608

